# Adrenal rather than central dysfunction limits HPA axis recovery after chronic glucocorticoid treatment in male mice

**DOI:** 10.1101/2025.04.30.651350

**Authors:** Lindsey S. Gaston, Brenna C. Jorgensen, Hannah R. Friedman, Marc S. Sherman, Joseph A. Majzoub

## Abstract

Glucocorticoid-induced adrenal insufficiency (GIAI) can persist for months after discontinuation of chronic corticosteroid therapy, placing patients at risk for life-threatening adrenal crises. This prolonged suppression has been attributed primarily to delayed restoration of hypothalamic–pituitary signaling based on indirect measures of central axis activity. To identify the rate-limiting site of hypothalamic-pituitary-adrenal (HPA) axis recovery, we evaluated the timing of functional and histologic recovery at each node of the axis following 8 weeks of dexamethasone (DEX) treatment in adult, male mice. DEX administration fully suppressed HPA axis activity. Unexpectedly, within one week of DEX withdrawal, hypothalamic *Crh* mRNA and plasma ACTH rebounded above control levels, whereas corticosterone (CORT) remained suppressed for an additional seven weeks. DEX-treated adrenals were markedly atrophic and contained large clusters of lipid-filled macrophages. Even after adjusting for macrophage content, CORT secretion was disproportionately low relative to the remaining adrenocortical cell mass despite supraphysiologic ACTH stimulation. The adrenal is thus the principal site of post-withdrawal GIAI, involving adrenocortical cell loss and a superimposed defect in steroidogenesis. We next tested whether preserving adrenal trophic signaling during glucocorticoid exposure could prevent GIAI. Adrenal function recovered more slowly in mice treated with DEX and daily cosyntropin (a synthetic ACTH analog) compared to those treated with DEX alone. In contrast, mice with non-suppressible endogenous ACTH due to targeted hypothalamic deletion of the glucocorticoid receptor maintained normal adrenal architecture and steroidogenic capacity despite prolonged DEX treatment. Pharmacologic treatments that mimic sustained trophic signaling to the adrenal during chronic glucocorticoid treatment may thus prevent GIAI.

## Introduction

Systemic glucocorticoids are prescribed to up to 3% of the population in developed countries for treatment of autoimmune and inflammatory diseases, malignancies, and to prevent allograft rejection^1,2^. Many additional patients are chronically treated with topical and inhaled formulations that can exert systemic effects even at moderate doses^3–5^. Like endogenous cortisol in humans and corticosterone in rodents, exogenous glucocorticoids engage the glucocorticoid receptor (GR) to suppress hypothalamic-pituitary-adrenal (HPA) axis activity through negative feedback. This feedback reduces the synthesis and secretion of corticotropin-releasing hormone (CRH) from the paraventricular nucleus of the hypothalamus and adrenocorticotropic hormone (ACTH) from the anterior pituitary^6^. In the absence of ACTH, a key trophic factor for the adrenal cortex, cortisol-producing cells of the zona fasciculata (zF) undergo both suppression of steroidogenesis and apoptotic cell loss^7,8^.

HPA axis suppression is typically transient in patients treated with short-term glucocorticoids. However, treatment exceeding four weeks can cause protracted adrenal insufficiency that can persist for up to a year after steroid withdrawal^9–11^. While awaiting HPA axis recovery, individuals are at risk for adrenal crisis with otherwise routine illnesses or trauma, which has been associated with longer hospitalizations, higher rates of ICU admissions, and even death in patients who are not promptly administered stress-dose steroids^12,13^.

Despite the ubiquity of glucocorticoid use and the morbidity of glucocorticoid-induced adrenal insufficiency (GIAI), there are currently no available treatments to prevent or hasten recovery from adrenal suppression. This is due, in part, to the prevailing view that prolonged GIAI is mediated by delayed, post-withdrawal recovery of the hypothalamus and pituitary, which are relatively inaccessible therapeutic targets. Post-withdrawal central suppression was first characterized by Graber et al., who reported that ACTH levels were inappropriately low to normal for at least two months in patients withdrawn from chronic glucocorticoids despite release from negative feedback^14^. Since direct ACTH assays had not yet been developed, systemic ACTH levels in this study were interpolated from the corticosterone responses of hypophysectomized rats injected with extracted patient plasma^14,15^. Another study cited inadequate ACTH responses to CRH stimulation testing in patients withdrawn from long-term prednisone as evidence of post-withdrawal central suppression. However, testing was conducted only 24 hours, or two biological half-lives, from the last prednisone dose, thus confounding glucocorticoid-mediated negative feedback and post-withdrawal suppression^16,17^. Post-withdrawal central suppression has also been described after surgical cure of Cushing’s syndrome, a state of endogenous glucocorticoid excess, though clinical studies have disagreed as to whether this is mediated by persistent hypothalamic or pituitary dysfunction^18–20^.

Interestingly, patients with congenital adrenal hyperplasia, who also require chronic, suppressive glucocorticoid therapy, exhibit elevated levels of ACTH at their morning hydrocortisone trough^21,22^. This suggests that some patients have rapid normalization of hypothalamic-pituitary function after release from glucocorticoid feedback inhibition. The mammalian HPA axis is highly conserved evolutionarily, and all rodent studies have reported rapid restoration of hypothalamic-pituitary function after glucocorticoid treatment. Basal and stimulated ACTH levels (following lipopolysaccharide, arecoline, and CRH administration) were recovered within days of discontinuing 5 or 7 days of methylprednisolone or dexamethasone, respectively, in rats^23,24^. In mice treated with dexamethasone for 30 days, plasma ACTH and corticotrope *Pomc* mRNA levels were elevated above control levels by 1-week post-withdrawal^25^. In all studies, corticosterone (CORT) levels and/or steroidogenic pathway gene expression remained low 5-7 days after discontinuing steroids despite normalization of adrenal weights^23–25^. Importantly, no animal studies to date have examined HPA axis recovery after longer courses of glucocorticoids known to cause protracted adrenal suppression in humans.

Given these discrepancies in the existing clinical and preclinical literature, we developed a murine model of GIAI to define the locus and mechanisms underlying persistent HPA axis dysfunction following withdrawal of long-term glucocorticoid treatment. After we discovered that adrenal dysfunction persists for months after steroid withdrawal despite rapid restoration of hypothalamic-pituitary signaling, we tested whether preserving adrenal trophic signaling during glucocorticoid exposure, using either a pharmacologic or genetic approach, could prevent GIAI.

## Materials and Methods

### Animals

All animal protocols were in accordance with the National Institutes of Health guidelines and were approved by the Animal Care and Use Committee of Boston Children’s Hospital (Protocol 00001983). Mice were housed on a 12-hour/12-hour light-dark cycle (lights on at 7 am) with ad libitum access to rodent chow and water unless otherwise indicated. Mice used for studies were all male and aged either 8 weeks at the time of DEX initiation (long-term studies) or 8-16 weeks (DEX-naïve controls or short-term studies).

***Wild-type (WT) mice:*** C57BL/6J mice (The Jackson Laboratory, Bar Harbor, ME) were maintained by breeding within our population.

***Sim1^Cre^:GR^fl/fl^ mice (HGRKO):*** Sim1^Cre^ mice (generous gift of Brad Lowell, Beth Israel Lahey Medical Center, Boston, MA) were bred to floxed GR exon 3 mice (B6.Cg-*Nr3c1^tm^*^1^*^.1Jda^*/J, Catalog #021021, The Jackson Laboratory, Bar Harbor, ME), both maintained on an inbred C57BL/6J background^26,27^. Littermates that were Cre negative but had GR exon 3 floxed alleles were used as controls. Genotyping was performed using the primers and conditions detailed in Table SI^28^.

***Pharmacologic treatments:*** Mice were treated with either water-soluble dexamethasone (Sigma-Aldrich 2915) at a concentration of 3.6 mcg/mL (∼20 mcg/day assuming an average water intake of 5.8 mL/day^29^) or no-drug control via drinking water for either 4 days (short-term, n=4-7) or 8 weeks (long-term studies, n=3-4 per timepoint for natural history study and n=6/group for rescue paradigms, *Fig. S1*^28^). When specified, mice were co-treated with 10 mcg/kg cosyntropin, (synthetic ACTH[1-24], Sandoz, NDC 0781-3440-95), dissolved at 10 mcg/mL in 0.9% saline. Cosyntropin was administered once daily by intraperitoneal injection at 16:30 to align with the murine circadian ACTH peak for either 4 days or 8 weeks, as above.

***Hormonal assays:*** All testing was performed, as specified, on separate days so that no mouse was bled more than once on a given day. Plasma for measurement of corticosterone and ACTH was obtained by rapid retroorbital phlebotomy without anesthesia into EDTA-coated capillary tubes with a total time from first handling the animal to completion of bleeding not exceeding 30 seconds. Basal/unstimulated phlebotomy was performed at 16:30 in a ventilated hood in the same room in which the animals were housed. We have previously shown that ACTH and CORT measurements obtained with this method are in the expected basal, unstimulated range^30,31^. Cosyntropin stimulation^32^, insulin-induced hypoglycemia testing (1 U/kg dose)^31^, and physical restraint (20 minutes)^31^ were all performed as previously published. Blood was collected on ice, and plasma was separated by centrifugation at 1500 RPM at 4°C and stored at −80°C until assay.

Plasma ACTH was measured by two-site sandwich ELISA (ALPCO, catalog #21-ACTHU-E01, RRID: <u>AB_3714847</u>). Plasma was diluted 1:10 with the ACTH 0 calibrator (ALPCO; 11-C-270-10-G25), and the assay was otherwise performed according to the manufacturer’s instructions. The assay sensitivity, defined as the smallest value distinguishable from zero at the 95% confidence limit, was 0.22 pg/mL; samples falling below this threshold were assigned a value of 0 pg/mL. Per manufacturer specifications, intra-assay CVs were 6.71% and 2.27% at mean concentrations of 42.2 and 269.9 pg/mL, respectively, and inter-assay CVs were 7.1% and 6.9% at mean concentrations of 42.3 and 287.8 pg/mL, respectively.

Plasma corticosterone was measured by competitive ELISA (ALPCO, catalog #55-CORMS-E01, <u>RRID: AB_3677658</u>) according to the manufacturer’s instructions. The assay sensitivity, defined as the lowest detectable level distinguishable from the zero calibrator at the 2SD confidence limit, was 6.1 ng/mL (or 0.61 mcg/dL); samples falling below this threshold were assigned a value of 0 mcg/dL. Per manufacturer specifications, intra-assay CVs were 8.9%, 7.3%, and 5.9% at mean concentrations of 62.8, 126.0, and 271.4 ng/mL, respectively, and inter-assay CVs were 7.2%, 8.2%, and 7.5% at mean concentrations of 59.3, 113.2, and 257.4 ng/mL, respectively.

***Necroscopy and tissue preparation:*** Animals were euthanized by rapid decapitation without anesthesia followed by necroscopy for isolation of hypothalami and adrenals. Thick coronal sections flanking the hypothalamus were prepared with a surgical blade and then fixed in 10% formalin overnight before transfer to 70% ethanol while awaiting paraffin embedding.

Adrenals were trimmed of surrounding fat tissue, rinsed in phosphate-buffered saline (PBS), and weighed before overnight fixation in 10% neutral buffered formalin.

### Single-molecule RNA in situ hybridization

Three 5 µm-thick FFPE coronal brain sections were mounted onto each slide, and every other slide was stained with hematoxylin and visualized with brightfield microscopy (Nikon E800). The paraventricular nucleus was identified histologically based on its characteristic “butterfly-shaped” appearance in coronal sections, which was used to anchor selection of 2 unstained slides immediately rostral and caudal to this structure for analysis. Single-molecule RNA in situ hybridization was performed using a catalog Mus musculus *Crh* probe (ACD #316091) and 2.5 HD RED detection kit (ACD #316091) following manufacturer’s protocol.

Slides were counter stained with 50% Gill’s Hematoxylin II and mounted with EcoMount. Positive control *Ppib (*ACD #313911) and negative control *dapB (*ACD #310043) were used in every experiment to ensure sample quality. Brightfield images at 40x magnification were generated with an Olympus VS120 Virtual Slide Scanner and then analyzed with semi-quantitative densitometry using the Weka Segmentation plugin for Fiji ImageJ per ACDBio instructions^14^. Analysis was restricted to freehand sections encompassing the characteristic “butterfly” morphology of PVH cell bodies. CRH expression, reported as arbitrary units (AU), was quantified by dividing the area of the *Crh* densitometric signal (µm^2^) by the total area of this region (mm^2^), normalized to this same value for dexamethasone naïve control animals.

Separately, chromogenic RNAScope was performed using the 2.5 HD RED detection kit following manufacturer’s instructions, as above, to detect *Cyp11b1* (Cat. #564431) and *Trem2* (Cat. #404111)

The RNAScope Multiplex Fluorescent V2 Assay kit was used according to manufacturer instructions for single-cell quantification of adrenal steroidogenic pathway genes. Probes were multiplexed across two panels, each hybridized on separate sequential sections. In Panel 1, *Star* (Cat. #543581) was assigned to channel C1 and detected with TSA-650, *Cyp11b1* (Cat. #564431) to C2 with TSA-520, and *Hsd3b2* (Cat. #467681) to C3 with TSA-570. In Panel 2, *Mc2r* (Cat. #318891) was assigned to C1 with TSA-650 and *Cyp11a1* (Cat. #809181) to C2 with TSA-520. All fluorescently labeled sections were counterstained with DAPI and incubated in TrueBlack, as above, prior to mounting. Slides were imaged on an Olympus VS200 slide scanner at 40x magnification using whole-slide scanning to capture at minimum an entire adrenal hemisphere per section.

### Total mRNA and protein staining

To quantify total mRNA *in situ*, FFPE sections were deparaffinized and antigen-retrieved in buffer TE (pH=9). We then adapted the protocol in Farny *et al.* and Hurt *et al.* to label polyA-capped RNAs on FFPE sections using Cy5-conjugated polyDT (26-4430-02, GeneLink)^33,34^.

Total protein *in situ* was measured using the N-Hydroxysuccinimide (NHS) ester method; NHS esters react with primary amines and well-approximate total protein mass^35,36^. Briefly, FFPE sections were deparaffinized and antigen-retrieved in buffer TE (pH=9). Slides were stained with NHS-Alexa 568 dye conjugates (Thermo Scientific A20003) in PBS for 30 minutes and then washed 5X in PBS.

### Paraffin section immunofluorescence

After fixation, adrenals were oriented with their broadest surface parallel to the cutting plane in 4% low-melting-temperature SeaPlaque agarose (Lonza, #50101). Agarose blocks were then dehydrated in ethanol, xylene, and embedded in paraffin blocks. Paraffin sections were cut at 5 µm thickness, and immunofluorescence was performed as described in Leng *et al.*^37^ with the antigen retrieval conditions and antibody concentrations detailed in Table SII^28^. When indicated, Streptavidin, Alexa Fluor 488 Conjugate (Invitrogen, #S112233) was added to the secondary antibody solution at a final concentration of 1:500. In all experiments, sections were incubated in TrueBlack Lipofuscin Autofluorescence Quencher (Biotium, #23007) in 70% ethanol at room temperature for 3 minutes and rinsed three times in PBS prior to mounting with Prolong Gold Antifade Mountant with DAPI (Invitrogen, #P36931). Images were acquired with either a Nikon upright Eclipse 90i microscope with a 20x/0.75 Plan-Apochromat objective or an Olympus VS120 Virtual Slide Microscope using a 20x objective and adjusted for brightness and contrast in ImageJ.

### Floating section immunofluorescence

After fixation, adrenals were oriented with their broadest surface parallel to the cutting plane in 4% low-melting-temperature SeaPlaque agarose (Lonza, #50101) and sectioned at a thickness of 100 µm with a vibratome. Individual slices were separated by free floating in PBS. Sections were wet mounted in PBS on Superfrost Plus slides (Fisher Scientific, 12-550-15) and imaged with a 4x objective on a Nikon E800 brightfield microscope. The area of each section was measured in ImageJ, and the largest section was selected for subsequent immunostaining. Floating section immunofluorescence was then performed as per Leng *et al*^37^ with the addition of Streptavidin, Alexa Fluor 488 Conjugate (1:500, Invitrogen, #S112233) to the secondary antibody solution. Images were acquired with a Nikon Ti2 microscope with a Yokogawa CSUW1 confocal unit, a Nikon Plan Apo Lambda 20x objective lens and a Zyla 4.2 PLUS sCMOS camera (ANDOR). All images were taken with identical laser strength, gain and exposure settings for the 405nm (used to capture autofluorescence), 488nm, and 647nm channels. Each image was acquired using the Large Image function in the NIS elements software. This function allowed us to take images in a 4×4 grid with 15% overlap, allowing the images to be stitched together. Brightness and contrast were adjusted in ImageJ.

### TUNEL Procedure

Apoptotic cells were detected using the Click-iT PLUS TUNEL Assay with Alexa 647 kit (Thermo Fisher C10619) according to manufacturer instructions, with the addition of TrueBlack (Biotium, #23007) prior to mounting, as above. Images were acquired with an Olympus VS120 Virtual Slide Microscope using a 20x objective and adjusted for brightness and contrast in ImageJ.

### Fluorescent Image Analysis

Fluorescent images were analyzed using QuPath (version 0.4.3), an open-source software for digital pathology.

***Positive Cell Detection:*** QuPath’s cell detection feature was employed to label cells as either Ki-67 or TUNEL positive based on co-localization of the 647 nm channel signal with the DAPI nuclear signal. Images were opened in fluorescence mode, and regions of interest (ROI) were manually defined to include only the adrenal cortex based on morphologic characteristics. ROI were further restricted to the zF for Ki-67 analysis, as defined by DAB2-negative cortical cells. The “cell detection” feature was then run specifying DAPI as the detection channel and the 647 nm channel nuclear (Ki-67 analysis) or cellular (TUNEL analysis) mean as the score compartment. Nuclear size and intensity threshold parameters for both were adjusted and inspected visually to ensure accurate positive cell detection. Any cell exhibiting signal overlap between the DAPI and 647 nm channels above the specified thresholds was classified as either TUNEL or Ki-67 positive and reported as a percentage among total cells.

***Morphometry:*** Morphometric analysis was performed on thick, floating section immunofluorescent specimens to quantify the area of either the adrenal cortex (Streptavidin staining, 488 nm channel), medulla (tyrosine hydroxylase immunostaining, 647 nm channel), or adrenal macrophages (autofluorescence, 408 nm channel) based on signal intensity across multiple channels using the “create thresholder” function in QuPath. Images were opened in fluorescence mode, and brightness/contrast of individual channels were adjusted for optimal signal visualization. ROI were manually defined to include the entire adrenal slice but not surrounding adipose tissue. For each channel, QuPath’s intensity thresholding was applied to ensure accurate delineation of stained or autofluorescent areas, as confirmed by visual inspection. As DEX treatment has been shown to attenuate streptavidin staining in the murine zF, we applied a lower threshold in 8+0 adrenals to ensure detection of all residual adrenocortical tissue^38^. The area of tissue with a signal above this threshold for each channel was classified as positive and reported in square microns.

**RNAScope Puncta Counting:** Fluorescent RNAScope puncta were quantified on a per cell level according a QuPath workflow outlined in an ACDBio technical note^39^ Raw multi-channel fluorescent images were uploaded to a new project in QuPath, and brightness/contrast was adjusted until individual puncta could be visualized with minimum background signal. For each image comprising an entire adrenal hemisphere, the zF was annotated using the closed polygon annotation tool based on cell morphology in the DAPI channel. Cell segmentation was performed using the “cell detection” function in the DAPI channel with default settings. Probe detection was then performed with the “subcellular detection” function. For each relevant channel, the detection threshold was adjusted until the software appropriately recognized puncta and clusters of puncta within the smallest possible area, with an expected spot size of 1 µm2, a minimum spot size of 0.5 µm2, and a maximum spot size of 2 µm2. Data were presented as the number of puncta per zF cell normalized to this same value for the dexamethasone naïve control condition (arbitrary units, or AU).

### Statistics

All statistical analyses were performed in GraphPad Prism v.10.3.1 (GraphPad Software, Boston, MA; www.graphpad.com). All experimental groups consisted of male same-cage littermates; group sizes of n = 3–5 reflect litter-based sampling. Formal a priori power calculations were not performed. Hormonal outcomes (stimulated ACTH and corticosterone) were analyzed as the principal endpoints; morphometric and histological analyses from the same animals were considered exploratory. Data are represented as means ± SEM.

Prior to parametric testing, normality was assessed using the Shapiro-Wilk test (Table SIII^28^). For multi-group comparisons intended for one-way ANOVA, homoscedasticity was additionally evaluated using the Brown-Forsythe test (Table SIII). For two-group comparisons, Welch’s unpaired t-test was used, which does not assume homogeneity of variance and therefore does not require prior assessment of homoscedasticity. When data for multi-group comparisons satisfied assumptions of normality and homoscedasticity, one-way ANOVA was performed with Bonferroni correction for post-hoc pairwise comparisons against the dexamethasone-naive (DN) control group. When normality assumptions were not met, data were analyzed non-parametrically with the Mann-Whitney U test for two-group comparisons and the Kruskal-Wallis test for multigroup comparisons using Dunn’s correction for pairwise, post-hoc comparisons vs DN controls. The only exception to this practice was in the analysis of ACTH recovery after long-term DEX. ACTH levels during long-term DEX treatment (8+0) showed a non-Gaussian distribution, as multiple values fell below the lower limit of assay detection. Rather than applying non-parametric testing to all groups, the 8+0 group was excluded from this analysis as the biological question concerned post-withdrawal recovery. The remaining groups (8+1 through 8+8 and DN controls) satisfied parametric assumptions and were analyzed by one-way ANOVA with Bonferroni correction for pairwise post-hoc comparisons against DN (Table SIII^28^). For continuous variables meeting normality assumptions (Shapiro-Wilk) but failing homoscedasticity (Brown-Forsythe), multi-group comparisons were performed using Welch’s one-way ANOVA with Dunnet’s T3 post-hoc testing for pairwise comparisons.

For morphometric analyses, only two adrenal glands were available from the 8+6 timepoint due to sample loss during tissue processing. These data are displayed in the corresponding histograms for transparency but were excluded from statistical comparisons due to insufficient sample size.

Nonlinear regression using an exponential growth equation was used to model stimulated corticosterone recovery (dependent variable) over time (independent variable); doubling time was computed as ln(2)/K, where K is the rate constant. When two exponential growth models were compared, the extra-sum-of-squares F-test was used to determine whether the models provided significantly different fits to the data. Homoscedasticity of regression residuals was confirmed by visual inspection of the residual plot (absolute residuals vs. fitted values), which showed no systematic increase in residual variance across the range of fitted values. A p value below 0.05 was considered statistically significant.

## Results

### Mice treated with long-term dexamethasone (DEX) have protracted adrenal insufficiency despite early hypothalamic-pituitary recovery

We first established when hormonal function recovered at each node of the HPA axis following long-term DEX treatment. Adult, male, WT mice were treated with either continuous DEX or vehicle (DEX-naïve, or DN) in the drinking water for 8 weeks. Starting at the time of DEX withdrawal (8+0) and at 1-2 week intervals through 8+8 (where A weeks of treatment plus B weeks of withdrawal = A+B), separate cohorts of mice underwent serial cosyntropin stimulation and insulin-induced hypoglycemia (IIH) testing followed by necroscopy (Model 1, Fig. S1^28^)

As expected, long-term DEX treatment suppressed *Crh* mRNA within the paraventricular nucleus of the hypothalamus (PVH; 0.15±0.04 in 8+0 vs. 1.0±0.1 arbitrary units [AU] in DEX-naïve (DN) control animals, p=0.0011). However, *Crh* expression recovered to control levels 1 week after DEX was discontinued (1.1±0.11 AU in 8+1 vs 1.0±0.1 in DN, p=0.96; Fig. 1*A, B*). Similarly, IIH-stimulated plasma ACTH concentrations were suppressed during the period of glucocorticoid exposure (4.2±3.6 pg/mL at 8+0) but rebounded significantly above control levels by 1-week post-withdrawal (1707±85 pg/mL at 8+1 vs. 875±44 pg/mL, p=0.0128). IIH-stimulated ACTH levels remained elevated through 4 weeks post-withdrawal and reached a peak two weeks after completion of DEX treatment (Fig. 1*C*). Despite this, both IIH- and cosyntropin-stimulated CORT levels remained significantly lower than DN controls until 8 weeks after DEX cessation (Fig. 1*D*, *E*). Thus, mice withdrawn from long-term DEX have delayed recovery of CORT secretion despite rapid restoration of hypothalamic-pituitary function.

**Figure 1.**
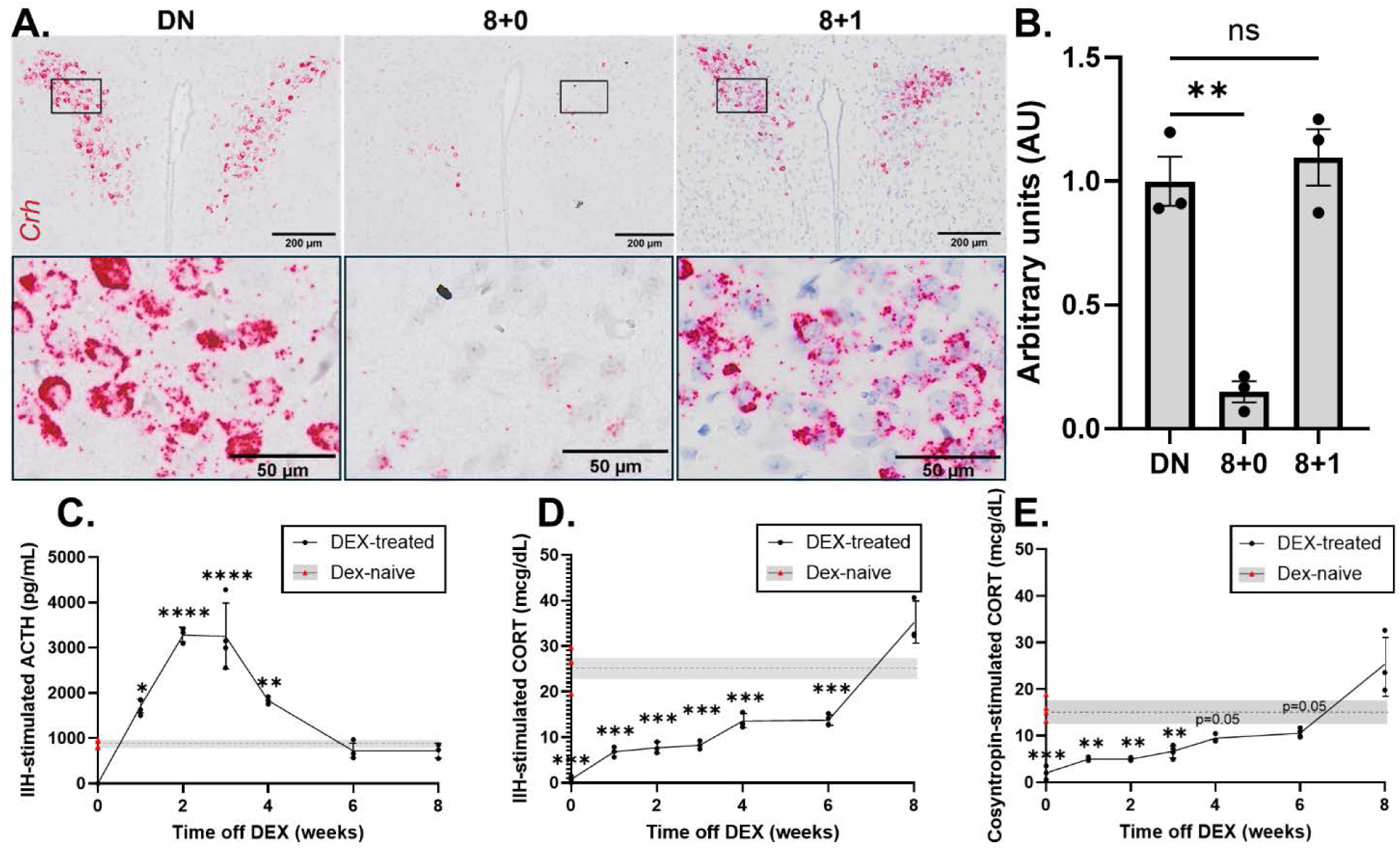
Corticosterone (CORT) secretion is suppressed for up to 8 weeks after discontinuing long-term DEX despite rapid hypothalamic-pituitary recovery. A) *In situ* hybridization (RNAScope) for basal *Crh* (red) in the paraventricular nucleus (PVH) of the hypothalamus in DEX-naïve (DN) animals and those treated with 8 weeks DEX both immediately prior to (8+0) and 1-week post-withdrawal (8+1; top row, scale bar=200 microns). Nuclei are labeled by hematoxylin (purple). Magnification of region within the black box is shown directly below (bottom row, scale bar=50 microns). B) Quantification of *Crh* mRNA expression from RNAScope images of the PVH from DN, 8+0, and 8+1 animals (n=3), normalized to DN controls. p=0.0006 by one-way ANOVA. Recovery of insulin-stimulated plasma C) ACTH (p<0.0001 by one-way ANOVA) and D) CORT (p<0.0001) and E) cosyntropin-stimulated CORT (p<0.0001) levels 0-8 weeks after withdrawal of an 8-week DEX course. Data shown as individual mice with mean (connecting line) ± SEM (error bars). Gray bar (mean± SEM) indicates the reference range for adult, male, wildtype, untreated (DEX-naïve or DN) controls (individual mice shown as red triangles overlying Y axis, n=4). * denotes p<0.05, ** p<0.01, *** p<0.001, and ****p<0.0001 for post-hoc pairwise comparisons with Bonferroni correction against DN.

### Long-term DEX induces adrenocortical atrophy with accumulation of large macrophage clusters

Having found the adrenal to be the limiting site of post-withdrawal GIAI, we then assessed adrenal mass and histology at select timepoints after chronic DEX. At 8+0, adrenal mass was reduced by 60% compared to controls (1.0±0.041 mg vs. 2.5±0.063 mg in DN, p<0.0001, Fig. 2*A*). Although adrenals from DEX-exposed mice weighed significantly less than those from DN controls through 6 weeks post-withdrawal, divergent patterns of recovery were noted for adrenal mass vs. function. Adrenal mass nearly doubled between 0 to 1 week post-withdrawal to 75% of controls (1.9±0.11 mg at 8+1) and remained relatively stable over the remaining recovery period. In contrast, CORT secretion was 75% lower than control values at 8+1 and more gradually recovered over the subsequent 7 weeks (Fig. S2*A*, *B*^28^; raw data derived from Figs. 1*D*, *E* and 2*A*) Hematoxylin and eosin staining revealed that DEX caused striking adrenocortical atrophy followed by gradual re-accumulation of cortical tissue in the 8 weeks post-withdrawal (Fig. 2*B*). Interestingly, DEX-exposed adrenals contained large, vacuolated structures with reduced nuclear density near the corticomedullary junction that persisted through 8 weeks post-withdrawal (Fig. 2*B*, magnification in 2*E* and S3*F*). These clusters were intensely autofluorescent, which was attenuated with the addition of TrueBlack, a lipofuscin autofluorescence quencher^40^ (Fig. S3*A*, *B*^28^). ISH with a 5’-Cy5-Oligod(T)30 demonstrated similar nuclear mRNA poly(A) content in these structures compared to surrounding cortical cells (Fig. S3*C*)^33,34^. The large, vacuolated, extranuclear space in these clusters was not strongly labeled with 568 NHS Ester (SE), a total protein stain (Fig. S3*D*)^28,35,36^. Together, this suggests that the multinucleated clusters observed in DEX-exposed adrenals contain live, lipofuscin-rich, biologically active cells rather than acellular, proteinaceous debris. These cells were negative for the steroidogenic enzyme *Cyp11b1* mRNA, a marker of zona fasciculata (zF) cells (Fig. S3*E*^28^).

**Figure 2.**
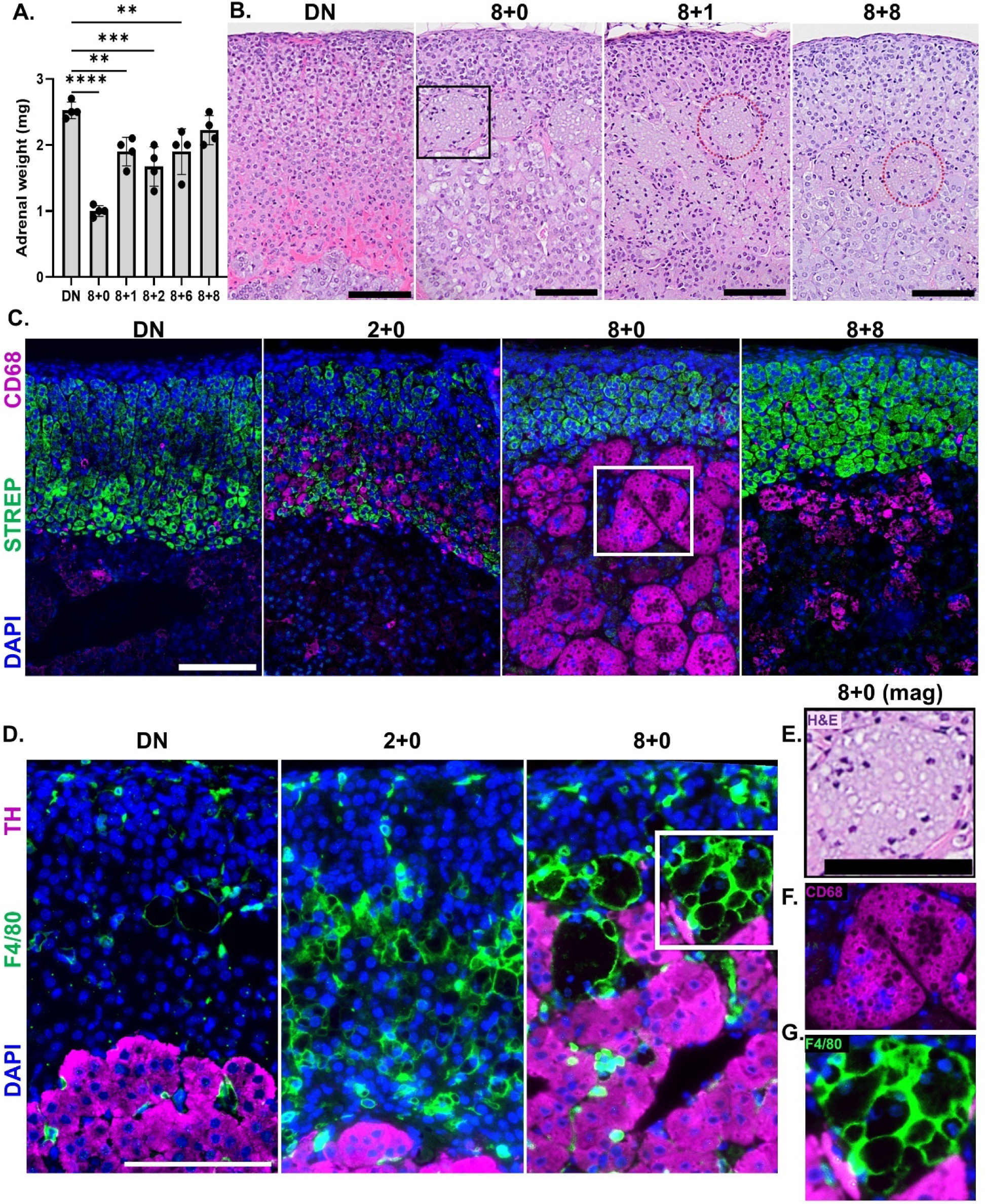
Long-term DEX induces adrenocortical atrophy with accumulation of large macrophage clusters that persist for at least 8 weeks post-withdrawal. A) DEX-treated adrenals weigh significantly less than DN controls through 6 weeks post-withdrawal. P<0.0001 by one-way ANOVA; * indicates p<0.05, ** p<0.01, *** p<0.001, and ****p<0.0001 for post hoc pairwise comparisons against DN with Bonferroni correction. B) Hematoxylin and eosin staining reveals large, poorly defined, vacuolated structures with decreased nuclear density at the corticomedullary junction in DEX-exposed adrenals that persist through 8-weeks post-withdrawal. Representative structures (outlined with red, dashed circles) correspond with clusters of cells that are positive for the classical macrophage markers C) CD68 and D) F4/80. C) Immunofluorescence for CD68 (magenta) with DAPI (blue) and 488-streptavidin (green) counterstains, the latter of which labels all biotin-rich cells of the adrenal cortex. Representative images shown for DN adrenals, those on DEX for 2-8 weeks (e.g. 2+0 to 8+0), as well as one (8+1) and eight (8+8) weeks after an 8-week DEX course. D) Immunofluorescence for the mouse macrophage marker F4/80 (green) and medullary marker tyrosine hydroxylase (TH; magenta) in DN and DEX-exposed adrenals (2+0 and 8+0). Magnification of representative macrophages in 8+0 glands from E) H&E and immunostaining for F) CD68 and G) F4/80. Scale bars=100 microns for all. All immunofluorescent staining performed with TrueBlack lipofuscin autofluorescence quencher (C, D, F, and G).

Immunostaining with the myeloid-specific marker, CD68, and macrophage-specific marker, F4/80, identified these cells as clusters of adrenal macrophages (AMs). Rare, mononuclear CD68- and F4/80-positive cells were visualized in the adrenal cortex (labeled with fluorescent 488-conjugated streptavidin^38^) of DN mice (Fig. 2*C*, *D*). Increased CD68- and F4/80-positive macrophages were present in the adrenal cortex after 2 weeks of DEX treatment (2+0). After 8 weeks of DEX (8+0), large AM clusters accumulated at the corticomedullary junction (Fig. 2*C*, *D*, magnified in 2*F*, *G*) and appeared to fragment and migrate into the adrenal medulla by 8 weeks post-withdrawal (8+8). AMs contained lipid, detected by both oil red O and BODIPY staining, but were negative for the lipid droplet associated proteins perilipin-1 and 2 (PLIN 1 and 2; Fig. S4*A*-*D*^28^). AMs were also positive for CD11c and *Trem2* by IF and ISH, respectively, both of which are expressed by lipid-associated macrophages in the context of metabolic-associated steatohepatitis and atherosclerosis (Fig. S4*E, F*)^28,41–44^.

### CORT secretion is disproportionately low relative to steroidogenic cell mass for at least 2 weeks post-DEX

As AMs likely contribute to adrenal mass but not biosynthetic function, we then asked specifically how the steroidogenic cell population contracted and recovered after chronic DEX treatment. To do this, we performed morphometric analysis on thick, immunostained adrenal sections, from which we selected the largest, equatorial one from each gland. Steroidogenic adrenocortical cells were labeled with 488-streptavidin^38^ (green), and the medulla was identified by immunofluorescence for tyrosine hydroxylase (red). Sections were not treated with TrueBlack so that AMs could be identified by endogenous autofluorescence (cyan; Fig. 3*A*).

**Figure 3.**
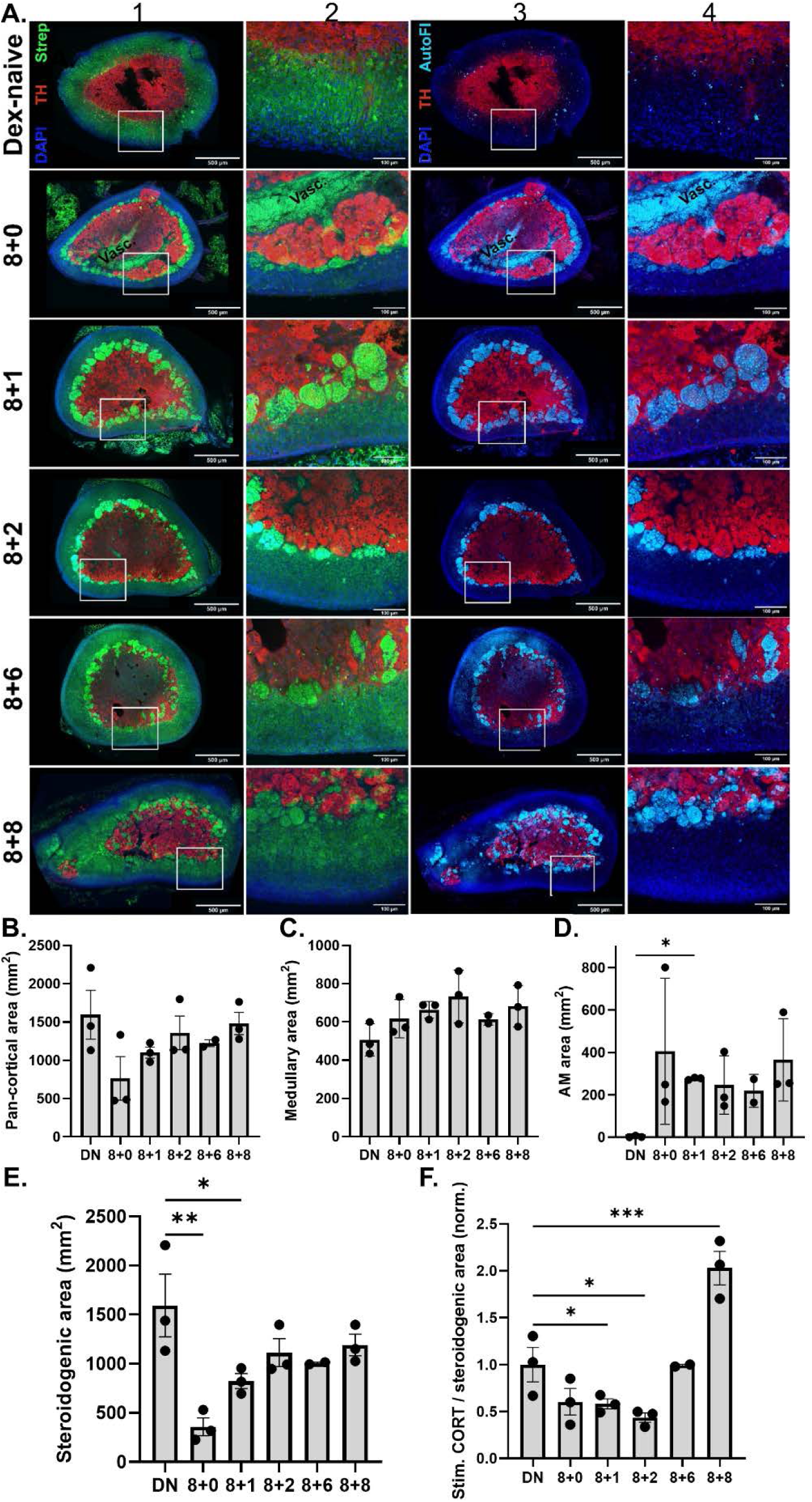
Stimulated CORT secretion is disproportionately low relative to adrenocortical cell mass for at least 2 weeks after withdrawal of long-term DEX. A) IF without TrueBlack for tyrosine hydroxylase (TH, red) DAPI (blue) and 488-streptavidin (green; columns 1 and 2) counterstains. Adrenal macrophages (AMs) were detected by autofluorescence (cyan; columns 3 and 4). Mice were either untreated (DN) or treated for 8 weeks DEX followed by withdrawal for 0-8 weeks (8+0, 8+1, 8+2, 8+6, and 8+8). Shown as 100 µM-thick, whole adrenal slices (columns 1 and 3, scale bar=500 microns) with magnification of the region in the white box (columns 2 and 4, scale bar=100 microns). Columns 1 and 2 imaged at wavelengths 405 nm, 488 nm, and 647 nm; columns 3 and 4 imaged at wavelengths 405 nm, 440 nm, and 647 nm. Morphometric analysis of B) pan-cortical area, which includes both streptavidin-positive and autofluorescent regions (p=0.26 by Kruskal-Wallis test; no significant post-hoc pairwise comparisons vs. DN using Dunn’s correction), C) Medullary area (regions (p=0.27 by Kruskal-Wallis test; no significant post-hoc pairwise comparisons vs. DN), D) Autofluorescent AM area (p=0.13 by Kruskal-Wallis test), E) steroidogenic cell type area, calculated as pan-cortical area minus AMs (p=0.0056 by one-way ANOVA). F) Ratio of steroidogenic cell type area to cosyntropin-stimulated CORT secretion, normalized to DN controls (p<0.0001 by one-way ANOVA). * p<0.05, ** p<0.01, *** p<0.001 for post hoc pairwise comparisons against DN using either Dunn’s (D) or Bonferroni correction (E-F).

As observed with adrenal weights (Fig. 2*A*), pan-cortical area, which includes both streptavidin-positive and autofluorescent regions, decreased with DEX treatment (764±284 mm^2^ in 8+0 vs. 1596±319.4 mm^2^ in DN) but nearly normalized by 1-week post-withdrawal (1100±72 mm^2^ in 8+1, Fig. 3*A*, *B*). Medullary (TH-positive) area was comparable at all time points, indicating that relatively equivalent, equatorial adrenal sections were used for morphometric analyses (p=0.027, Fig. 3*C*). Autofluorescent AM area increased with DEX treatment and remained stably elevated through 8 weeks post-withdrawal (Fig. 3*A*, *D*). Steroidogenic cell type area, computed by subtracting AMs (cyan autofluorescence) from the pan-cortical (green) area, markedly decreased with DEX treatment (272±47 mm^2^ in 8+0 vs. 1286±153 mm^2^ in DN, p=0.0019) but more than tripled in the first week post-withdrawal (887±70mm^2^; Fig. 3*A*, *E*).

To examine how adrenal function relates to adrenocortical cell mass, we calculated the ratio of cosyntropin-stimulated CORT secretion to the steroidogenic cell type area, normalized to DN controls (Fig. 3*F*). This ratio decreased at the end of DEX treatment (0.60±0.14, p=0.05) but reached its lowest values at 1- and 2-weeks post-DEX (0.58±0.05 and 0.44±0.05, p=0.04 and 0.01 vs. DN). By 8+6, the ratio had returned to baseline (0.99±0.01) and was elevated relative to DN by 8+8 (2.0±0.18, p=0.0001 vs DN). These findings indicate that, for at least 2 weeks after DEX withdrawal, CORT secretion is disproportionately low compared to the size of the non-macrophage adrenal cortex, even under supraphysiologic ACTH stimulation.

Immunofluorescence for CD31 revealed preserved vascularity in both DEX-treated (8+0) and withdrawn (8+1) adrenals (Fig. S5*A*^28^). The zona glomerulosa, labeled by Dab2, thickened during chronic DEX exposure (45±2.6 µm at 8+0 vs. 36 ± 0.96 µm in DN, p=0.04) but returned to baseline by 8+1 (36±2.2 µm, p>0.99; Fig. S5*B*, *C*^28^). Ki-67 staining demonstrated a robust increase in adrenocortical proliferation within one week of DEX withdrawal (17±1.5% of 8+1 cells vs. 2.0 ± 1.0% of DN cells, p=0.0001; Fig. S5*B* and *D*^28^), consistent with the known mitogenic effects of ACTH^45^. Together, these findings indicate that disproportionately low CORT secretion relative to steroidogenic cell type area cannot be explained by impaired adrenal vascularity, zonation, or trophic signaling but instead reflects a selective defect in CORT biosynthesis or secretion.

### Star mRNA expression per zF cell is incompletely restored after DEX withdrawal despite supraphysiologic ACTH stimulation

We next asked whether this defect in CORT secretion could be explained by impaired transcriptional regulation of steroidogenic genes in response to an acute stressor. Adrenals from dexamethasone-naïve (DN), DEX-treated (8+0), and DEX-withdrawn (8+1) animals were harvested 2 hours after insulin-induced hypoglycemia testing (Fig. 1*C-D*). mRNA content was analyzed by multiplexed single-molecule fluorescent in situ hybridization (RNAScope) to distinguish downregulation of steroidogenic gene expression at the single-cell level from reduction in zona fasciculata cell number (Fig. 4*A-E*). Data are expressed as mRNA puncta per zF cell, normalized to DN controls (Fig. 4*F-J*).

**Figure 4.**
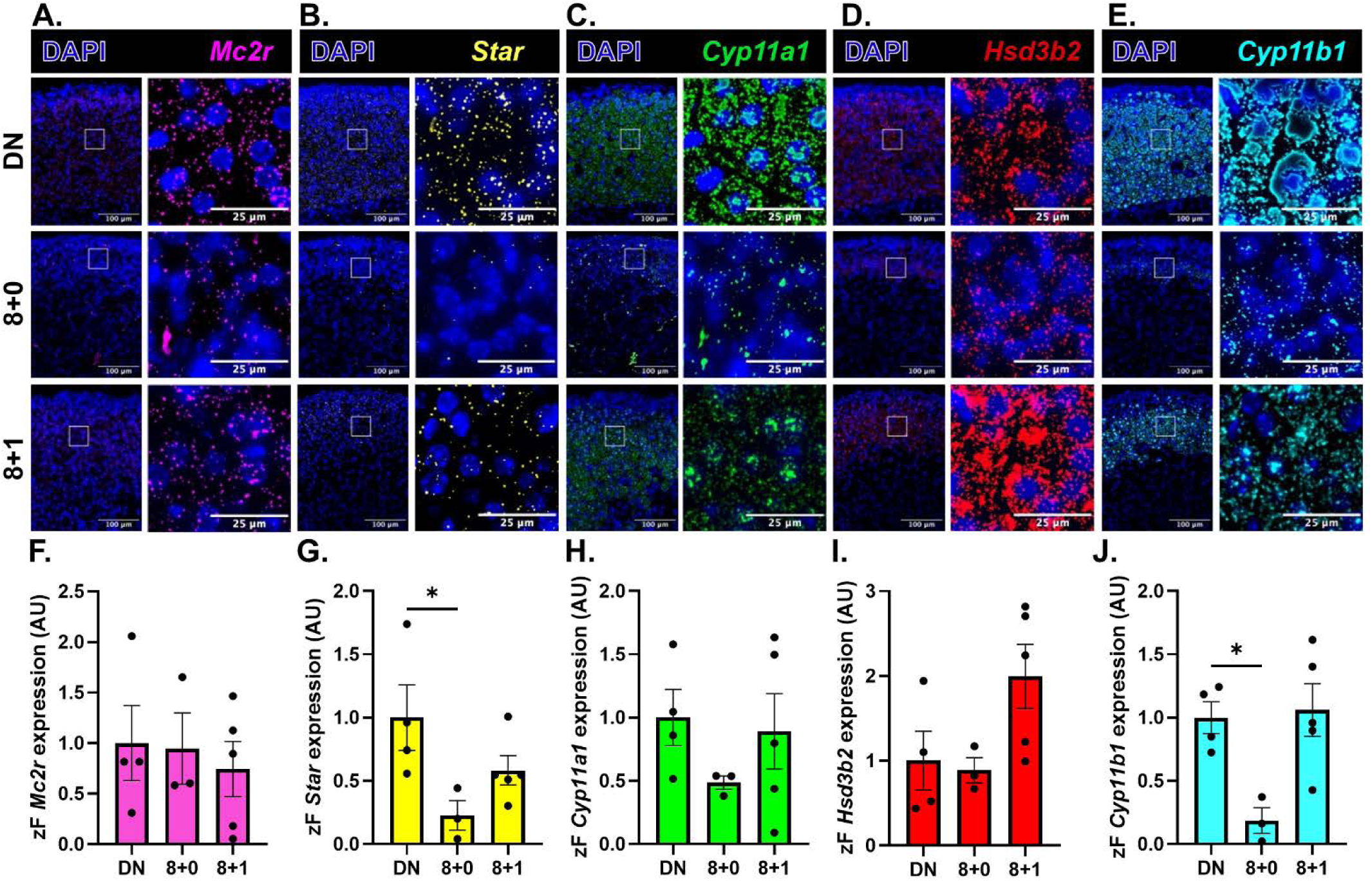
Steroidogenic pathway gene expression per zona fasciculata cell in dexamethasone-naïve (DN), dexamethasone-treated (8+0), and dexamethasone-withdrawn (8+1) adrenals in response to an acute stressor. Adrenals were harvested 2 hours after insulin administration (1 u/kg IP). Representative images obtained at 40x magnification from multiplexed fluorescent *in situ* hybridization (RNAScope, Multiplex Fluorescent V2 Assay, ACDBio) for A) *Mc2r* (magenta), B) *Star*, C) *Cyp11a1,* D *Hsd3b2*, and E) *Cyp11b1.* Data shown as widefield view encompassing all adrenal zones (left, scale bar=100 microns) with magnification of the representative zF region enclosed within the white box (right, scale bar=25 microns) to allow visualization of individual puncta. Quantification of mRNA puncta per zF cell normalized to DN controls: F) *Mc2r:* p=0.9268 by Kruskal-Wallis test; no significant post-hoc pairwise comparisons vs. DN using Dunn’s correction, G) *Star* (p=0.056 by one-way ANOVA), H) *Cyp11a1* (p=0.44 by Kruskal-Wallis test; no significant post-hoc pairwise comparisons vs. DN), I) *Hsd3b2* (p=0.086 by one-way ANOVA; no significant pairwise comparisons vs. DN), J) *Cyp11b1* (p=0.0175 by one-way ANOVA). * indicates p<0.05 for post-hoc pairwise comparisons against DN with Bonferroni correction.

Expression of *Mc2r*, encoding the ACTH receptor, did not differ significantly across treatment conditions (p = 0.9268 by Kruskal-Wallis; p > 0.9999 for both 8+0 and 8+1 vs. DN; Fig. 4*A, F*). Per-cell expression of multiple steroidogenic enzyme transcripts was reduced during long-term dexamethasone treatment (8+0): *Star* (0.23 ± 0.12 vs. 1.0 ± 0.26 AU in DN; p = 0.04; Fig. 4*B, G*), *Cyp11a1* (0.49 ± 0.052 vs. 1.0 ± 0.22; p = 0.36; Fig. 4C*, H*), and *Cyp11b1* (0.19 ± 0.10 vs. 1.0 ± 0.13; p = 0.03; Fig. 4*E, J*). By one-week post-withdrawal (8+1), *Cyp11a1* and *Cyp11b1* expression had returned to control levels (0.89 ± 0.30 and 1.06 ± 0.21, respectively; p > 0.9999 for both vs. DN, Fig. 4*C, E, H,* and J). In contrast, *Star* mRNA content per zF cell remained below control levels at 8+1 (0.58 ± 0.11 vs. 1.0 ± 0.26; p = 0.12, Fig. 4B*, G*) despite exposure to 2-fold higher plasma ACTH concentrations. *Hsd3b2,* another positively ACTH-regulated transcript, was instead upregulated two-fold in 8+1 adrenals compared to DN (2.0 ± 0.38 vs. 1.0 ± 0.35; p = 0.13, Fig. 4*D, I*). Together, these data raise the possibility of selectively impaired induction of *Star*, the rate-limiting step in steroidogenesis, during an acute stressor despite supraphysiologic ACTH stimulation.

### Maintenance of endogenous ACTH secretion but not intermittent cosyntropin administration during the period of DEX exposure prevents GIAI

Having localized post-withdrawal GIAI to the adrenal, we next tested whether maintaining trophic stimulation during glucocorticoid exposure could prevent adrenal suppression. Wild-type mice were treated with DEX as above and received daily, intraperitoneal injections of cosyntropin (ACTH[1-24]; DEX+COS) for either 4 days (short-term) or 8 weeks (long-term). Long-term cohorts underwent either a) serial cosyntropin stimulation testing at 1–2-week intervals from 0-8 weeks after withdrawal, or b) necroscopy at 8+1 (Model 2, Fig. S1). Short-term DEX+COS treatment significantly but variably reduced DEX-induced adrenocortical apoptosis (5.7±1.7% vs. 11±0.8% in 4D DEX glands, p=0.045; Fig. 5*A*, *B*). In contrast, after 8 weeks of treatment (8+0), cosyntropin-stimulated CORT levels remained equally suppressed in DEX+COS and DEX-only mice (2.0±1.0 µg/dL vs. 2.0±0.61 µg/dL, p=0.99). Following withdrawal, CORT recovery was significantly delayed in DEX+COS mice (doubling time=5.2 weeks) compared to DEX-only mice (2.5 weeks; p=0.0009, Fig. 5*C*). Morphometric analysis at 8+1 revealed comparable steroidogenic cell type area between DEX+COS and DEX-only adrenals (0.49±0.13 and 0.51±0.046, p=0.88, expressed as normalization to DN) but reduced AM area in the DEX+COS group (28±8.1 vs. 82±0.95, p=0.1, Fig. *5D-G*). AM morphology was also heterogeneous, ranging from scattered smaller cells throughout the zona fasciculata to typical aggregates at the corticomedullary junction (Fig. 5*D*). Thus, although intermittent pharmacologic ACTH stimulation partially limits adrenocortical apoptosis and macrophage accumulation, it fails to sustain steroidogenic capacity and ultimately slows functional recovery.

**Figure 5.**
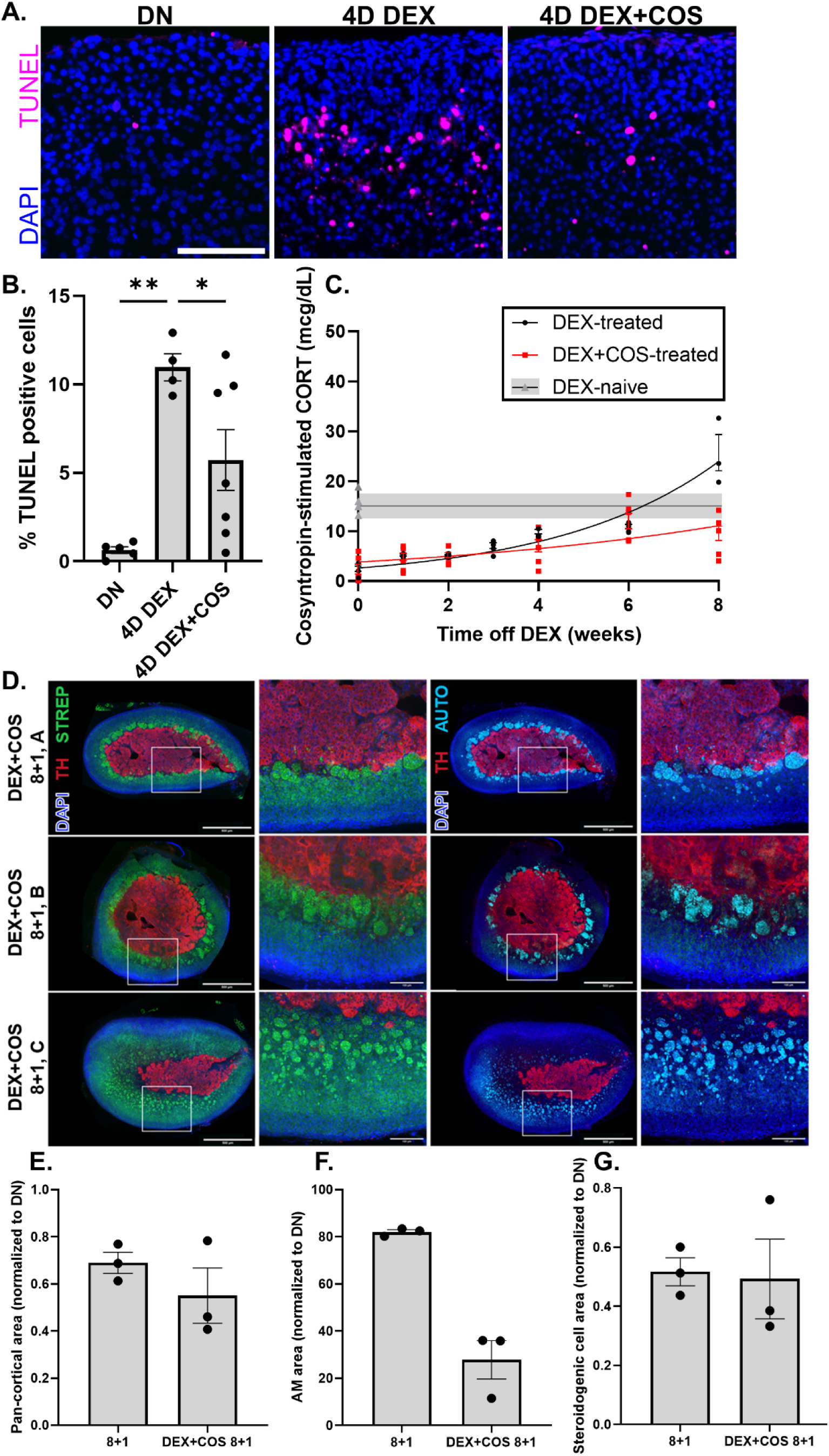
Combined dexamethasone and daily, intraperitoneal cosyntropin (DEX+COS) treatment partially attenuates cortical apoptosis and macrophage accumulation but fails to preserve adrenal function. A) Terminal deoxynucleotidyl transferase dUTP nick end labeling (TUNEL, magenta) in adrenals from WT mice treated for 4 days with vehicle (DN), DEX alone (4D DEX), or DEX with daily, intraperitoneal cosyntropin (4D DEX+COS), scale bar=100 microns with B) quantitation. p<0.0001 by Welch’s ANOVA; p=0.0017 for 4D DEX vs. DN and p=0.045 for DEX+COS vs. 4D DEX using Dunnet’s T3 post-hoc testing for pairwise comparisons. DN and 4D DEX data also shown in Fig. 7A. C) After withdrawal, cosyntropin-stimulated CORT secretion recovers more slowly in mice treated for 8 weeks with DEX+COS (red) vs. DEX alone (black). Data shown as individual mice with error bars centered on mean value ± SEM. Gray bar (mean ± SEM) indicates the reference range for adult, male, wildtype, DEX-naïve controls. Stimulated CORT recovery was modeled with exponential functions yielding doubling times of 5.2 weeks after DEX+COS (*Y* = 3.8 ∗ *e^0.13X^* R^2^=0.4, red curve) vs. 2.5 weeks after DEX (*Y* = 2.6 ∗ *e^0.28X^*; R^2^=0.9, black curve). p= 0.0009 by extra sum-of-squares F test. D)Thick, floating section IF without TrueBlack for tyrosine hydroxylase (TH, red), with DAPI (blue), and 488-streptavidin (green) in adrenals withdrawn from long-term DEX+COS (8+1); methods and visualization as per Figs. 3 and 4. All biological replicates are shown given significant morphologic heterogeneity among animals. Morphometric analyses of E) pan-cortical (p=0.36 by Welch’s t-test), F) AM (p=0.1 by Mann-Whitney U test), and G) steroidogenic cell type area (p=0.87 by Welch’s t-test, computed as per Figs. 3 and 4) in 8+1 DEX-monotherapy and DEX+COS glands, all normalized to DN controls. Measurements from DN and 8+1 DEX-only adrenals previously shown in Fig. 3.

We then tested whether sustained, endogenous ACTH signaling during the period of glucocorticoid exposure prevents GIAI (Model 3, Fig. S1). We generated Sim1^Cre^:GR^fl/fl^ mice (HGRKO), which harbor selective hypothalamic deletion of exon 3 of *Nr3c1,* encoding the glucocorticoid receptor (GR). These mice were previously shown by Laryea *et al.* to have 3-fold higher levels of *Crh* mRNA within the PVH compared to wild-type (WT) littermate controls, which was associated with ACTH and CORT levels that maintained circadian rhythmicity but were significantly higher in the basal state at the circadian peak and nadir as well as in response to restraint stress^27^. They further demonstrated that CORT levels decreased by 50% in WT controls after overnight dexamethasone administration but paradoxically increased in HGRKO mice^27^. These results identify HGRKO mice as having increased HPA axis activity that is resistant to glucocorticoid feedback inhibition, prompting us to ask whether these mice were similarly protected against DEX-induced adrenocortical apoptosis and functional suppression after chronic glucocorticoid treatment.

HGRKO mice or Cre-negative littermate controls (HGRWT) were treated with DEX for either 4 days or 8 weeks. HPA axis activity was assessed pre-treatment and on chronic DEX (8+0) in the unstressed state, following physical restraint, and after cosyntropin stimulation. On long-term DEX treatment (8+0), GR immunoreactivity was present in 94% of PVH neurons of HGRWT mice vs. 23% in HGRKO mice (Fig. 6*A*, *B*). *Crh* expression in the PVH was suppressed in these same DEX-treated HGRWT (49±12 AU) but not HGRKO animals (244±2.2 AU; Fig. 6*C, D*). GR was thus mostly deleted from the PVH, which conferred resistance to DEX-induced feedback inhibition of *Crh* mRNA expression.

**Figure 6.**
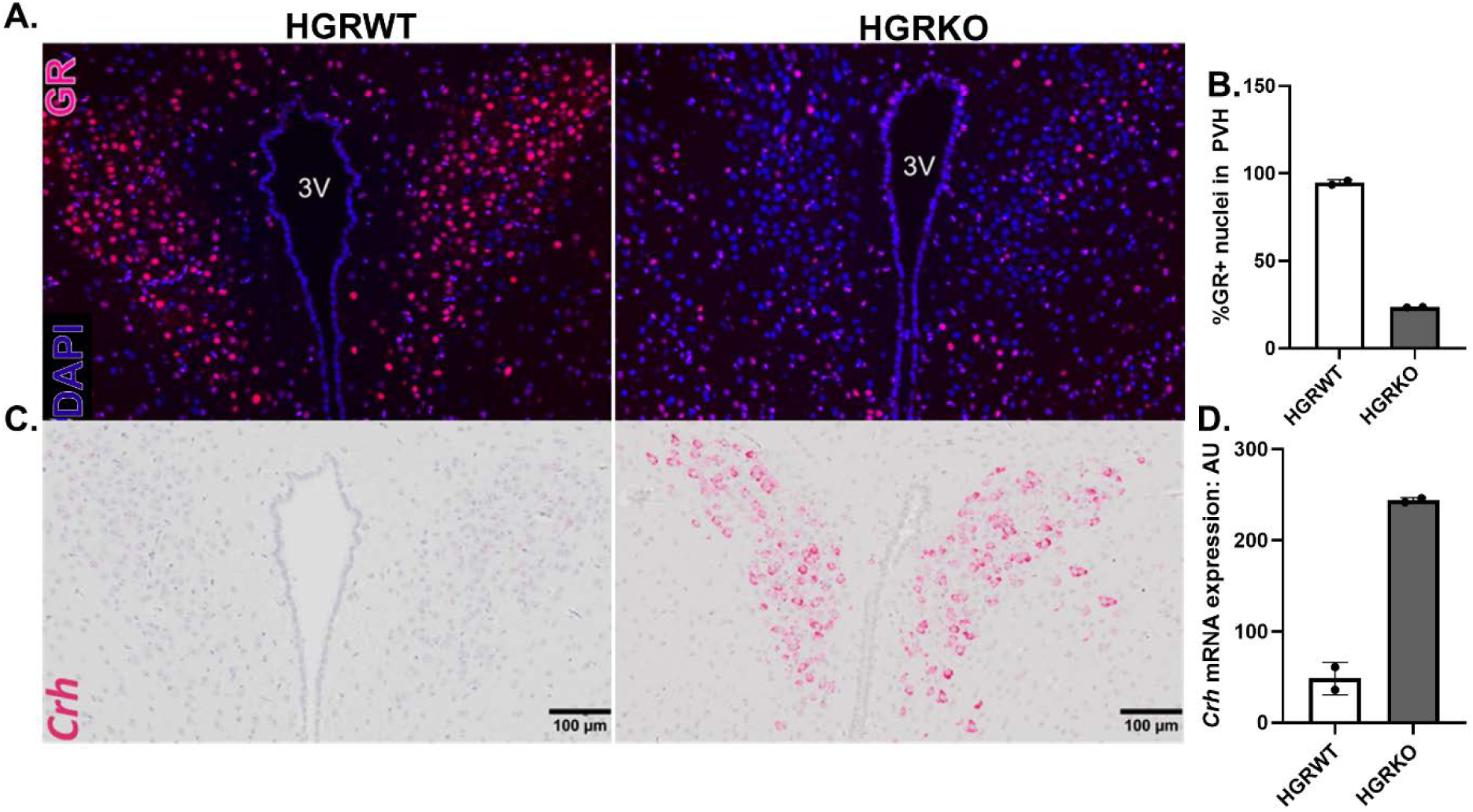
Selective deletion of the glucocorticoid receptor from hypothalamic neurons protects against glucocorticoid-mediated suppression of *Crh* expression. A) Immunofluorescence for the glucocorticoid receptor (GR, red; DAPI, blue) in the paraventricular nucleus of the hypothalamus (PVH) of DEX-treated HGRWT (Cre-negative controls; left panel) and HGRKO (Sim1^Cre+^:GR^fl/fl^; right panel) mice with B) quantification of the percentage of GR-positive nuclei. C) *In situ* hybridization (RNAScope) for *Crh* (red) in PVH sections from DEX-treated HGRWT and HGRKO animals with D) quantification. Scale bars=100 microns.

Apoptosis was uniformly rare in DN (0.63±0.19%) and short-term DEX-treated HGRKO glands (1.7±0.9%, p=0.61), compared to the increase found in DEX-treated WT glands (11±0.8%, p=0.0023 vs. DN and p=0.0006 vs. 4D DEX HGRKO; Fig. 7A, *B*). HGRKO mice had higher basal ACTH concentrations than HGRWT controls (470±76 vs. 104±55 pg/mL, *p*<0.0079) that were not suppressed on long-term DEX (385±64 vs. 0±0 pg/mL; *p*<0.0238; Fig. 7*C*). Pre-treatment basal and stimulated CORT levels were expectedly higher in HGRKO compared to HGRWT mice (Fig. 7*D*). On long-term DEX, basal, restraint- and cosyntropin-stimulated CORT were all markedly suppressed in HGRWT mice, whereas these levels remained high in HGRKO animals (basal: 1.5±0.98 vs. 14±1.6 µg/dL, p=0.0003; restraint: 1.9±1.6 vs. 33±4.5 µg/dL, p=0.0005; cosyntropin-stimulated: 1.1±0.62 vs. 33±3.9 µg/d, p=0.024, Fig. 7*D*).

**Figure 7.**
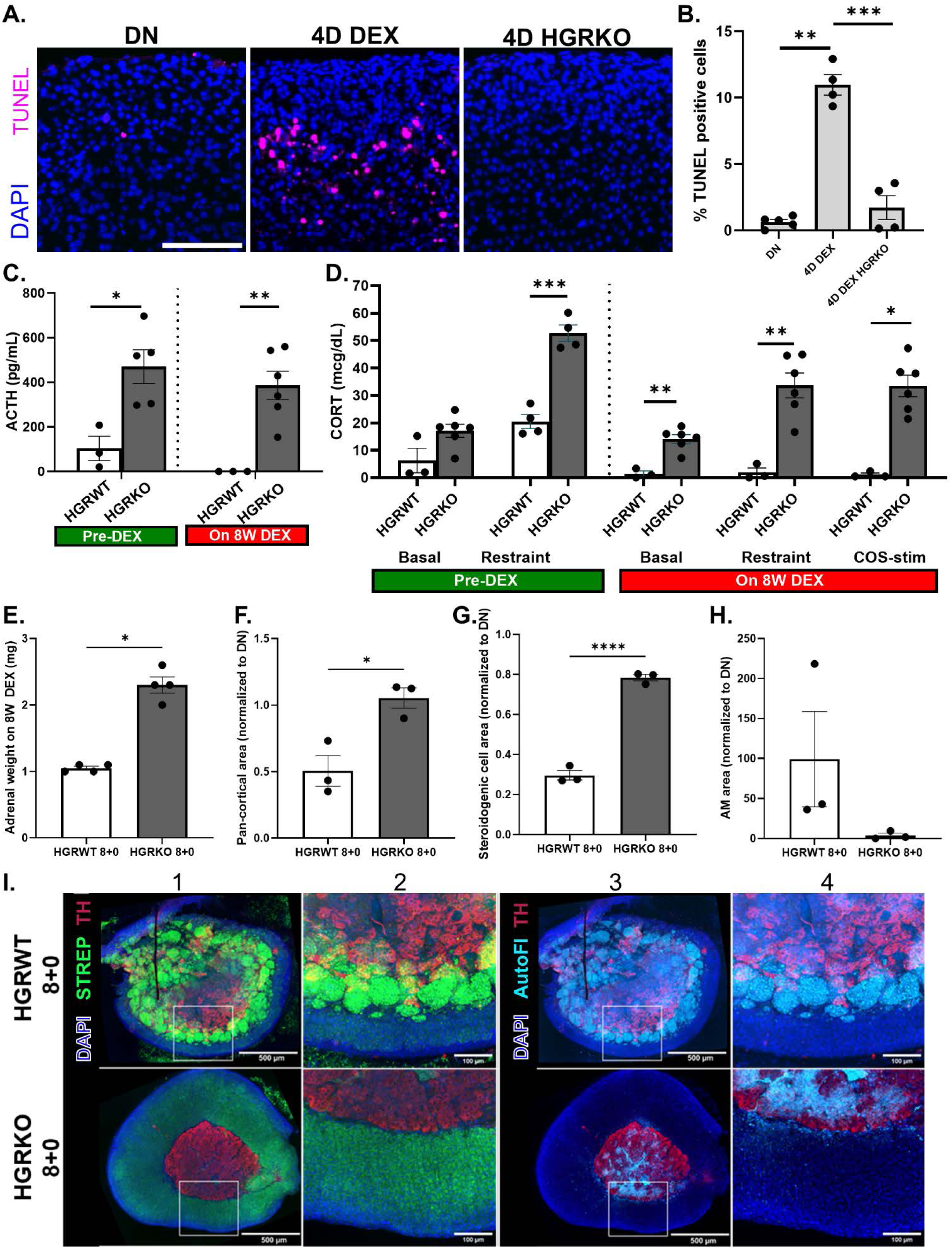
Selective deletion of the hypothalamic glucocorticoid receptor (GR) confers protection against DEX-induced adrenal suppression. A) Short-term (4 days) DEX induces apoptosis in wild-type (4D DEX) but not Sim1^Cre^:GR^fl/fl^ (4D HGRKO) mice, as shown by TUNEL staining (magenta), scale bar=100 microns with B) quantification. p<0.0001 by Welch’s ANOVA, **p<0.01 and *** p<0.001 by Dunnet’s T3 post-hoc testing for pairwise comparisons. HGRKO mice and Cre-negative, functionally wild-type littermate controls (HGRWT) mice were then treated with DEX for 8 weeks. Hormonal function was assessed pre-treatment and on long-term DEX. C) Basal (unstressed circadian peak) ACTH levels in HGRWT vs. HGRKO animals prior to DEX (left of dashed line, p=0.0079 by Welch’s t-test) and on 8 weeks of DEX (right of dashed line, p=0.0238 by Mann-Whitney U test). D) CORT secretion in the basal state (p=0.09 by Mann-Whitney U test for pre-DEX; p=0.0003 by Welch’s t-test for on 8W DEX) , following 20 minutes of physical restraint (Restraint; p=0.0002 for pre-DEX and p=0.0005 for on 8W DEX, both by Welch’s t-test), and following cosyntropin stimulation (COS-stim., p=0.024 for on 8W DEX by Mann-Whitney U test) in HGRWT vs HGRKO animals, either prior to DEX treatment (left of dashed line) vs. on 8 weeks of DEX (right of dashed line). E) Adrenal weights (p=0.029 by Mann-Whitney U test) and morphometric analyses for F) pan-cortical area (p=0.017 by Welch’s t-test), G) steroidogenic cell type area (p<0.0001 by Welch’s t-test), and G) AM area (p=0.1 by Mann-Whitney U test) in HGRWT and HGRKO adrenals on long-term DEX (8+0), all computed as per Fig. 3 and normalized to DN adrenals from Fig. 3. I) Representative thick, floating section IF without TrueBlack for tyrosine hydroxylase (TH, red), with DAPI (blue), and 488-streptavidin (green); methods and visualization as per Fig. 3. * p<0.05, ** p<0.01, *** p<0.001, **** p<0.0001 by Mann-Whitney U-test or Welch’s t-test, as indicated for HGRWT vs. HGRKO, C-H).

Adrenal mass (2.3±0.12 in HGRKO vs. 1.05±0.03 mg in HGRWT, p=0.029), pan-cortical area (1.1±0.077 in HGRKO vs. 0.51±0.12 in HGRWT, p=0.017, expressed as normalization to DN), and steroidogenic cell type area (0.78±0.016 in HGRKO vs. 0.30±0.024 in HGRWT, p<0.0001) were all significantly higher in HGRKO animals compared to controls (Fig. 7*E-G*). HGRKO adrenals were morphologically normal and lacked the large clusters of adrenal macrophages observed in HGRWT controls (AM area=3.7±2.9 in HGRKO vs. 99±60 in HGRWT, p=0.01, Fig. 7*H* and *I*). Sustained, CRH-driven, endogenous ACTH signaling during long-term DEX treatment thus prevents GIAI and cortical atrophy.

### Mc2r expression does not differ among DEX-treated WT, DEX+COS-treated WT, and DEX-treated HGRKO mice

ACTH transcriptionally regulates expression of its own receptor, *Mc2r,* in a positive, dose- and duration-dependent manner^46,47^. We therefore asked whether the failure of intermittent cosyntropin to prevent DEX-induced adrenal suppression reflected impaired regulation of the receptor through which it signals. RNAScope for *Mc2r* was performed on adrenals from WT mice that were either dexamethasone unexposed (DN, controls) or treated for 4 days with DEX either as monotherapy (WT 4D DEX) or in combination with cosyntropin (WT 4D DEX+COS), as well as HGRKO mice treated for 4 days with DEX (HGRKO 4D DEX, Fig. 8*A*). Within the zona glomerulosa (zG), *Mc2r* mRNA expression was similar among treatment conditions (Fig. 8*B*). Within the zF, per-cell *Mc2r* mRNA was reduced in all DEX-exposed groups relative to DN (1.0±0.37 AU): WT 4D DEX (0.21±0.06, p=0.041), WT 4D DEX+COS (0.20±0.08, p=.0.038), and 4D DEX HGRKO (0.16±0.053, p=0.029, Fig. 8*C-D*). Cosyntropin co-treatment thus fails to rescue *Mc2r* expression in DEX-treated WT mice. However, HGRKO mice exhibit equivalently suppressed *Mc2r* mRNA yet remain resistant to GIAI, indicating that *Mc2r* downregulation is not sufficient to cause DEX-induced adrenal suppression.

**Figure 8.**
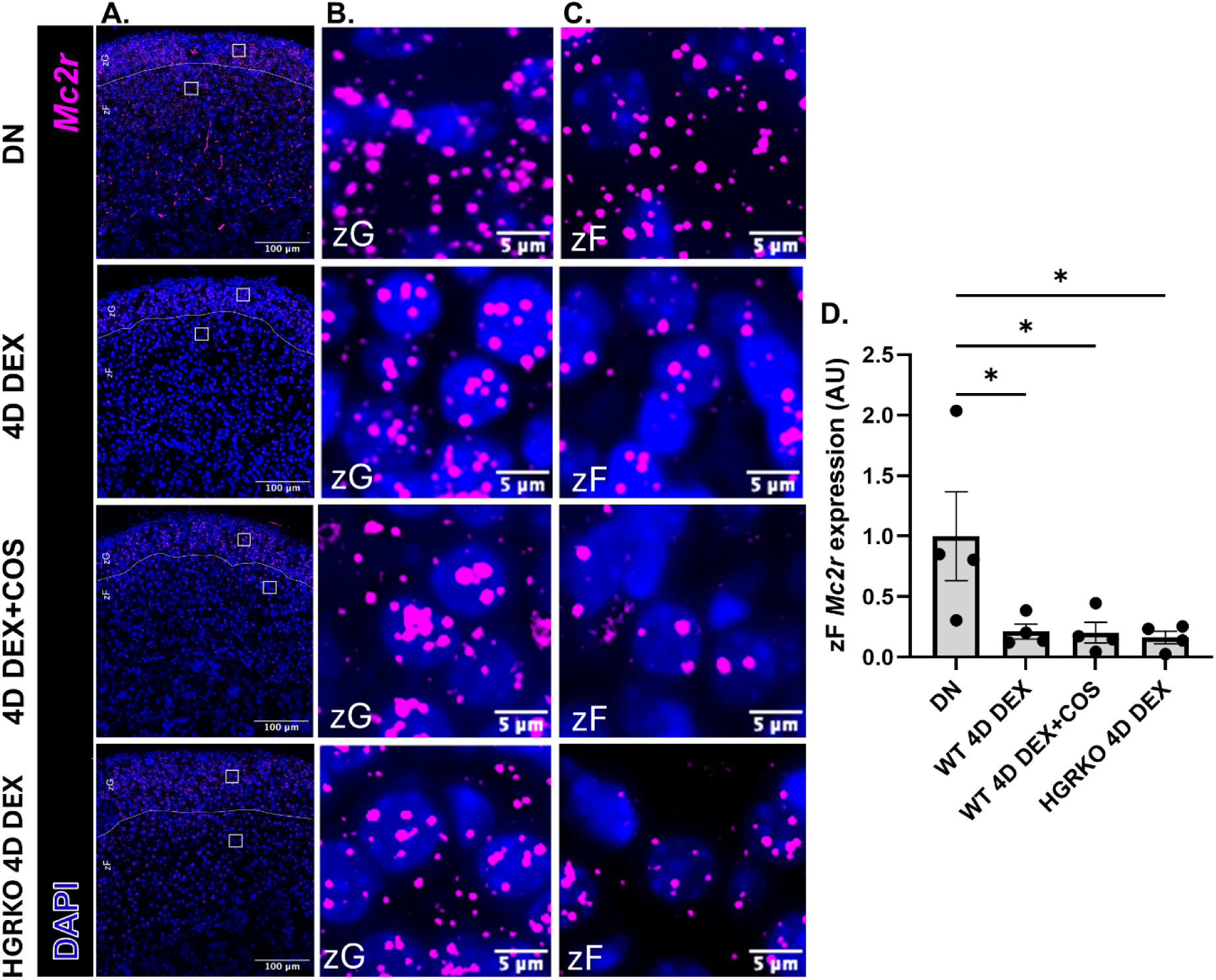
*Mc2r* mRNA content per zF cell does not differ among DEX-treated WT, DEX+cosyntropin-treated WT, and DEX-treated HGRKO mice. WT mice were either dexamethasone unexposed (DN, controls) or treated for 4 days with DEX either as monotherapy (4D DEX) or in combination with cosyntropin (4D DEX+COS). HGRKO mice were treated for 4 days with DEX (HGRKO 4D DEX). Representative images obtained at 40x magnification from fluorescent *in situ* hybridization (RNAScope, Multiplex Fluorescent V2 Assay, ACDBio) for *Mc2r.* Data shown as A) widefield views encompassing all adrenal zones (scale bar=100 microns), B) magnification of the boxed zG region, showing grossly similar *Mc2r* expression in this layer across treatment conditions, and C) magnification of the boxed zF region (scale bar=5 microns for *B* and C) with D) quantification of *Mc2r* puncta per zF cell normalized to DN controls (p=0.0263 by one-way ANOVA). *p<0.05 by post-hoc pairwise testing vs. DN with Bonferroni correction.

## Discussion

Using a murine model of GIAI, we found that hypothalamic-pituitary function rapidly recovered after cessation of chronic glucocorticoids, whereas adrenal suppression persisted for up to 8 weeks. DEX-exposed adrenals exhibited marked cortical atrophy and infiltration with large, lipid-laden macrophages. Even after accounting for these macrophages, CORT production was disproportionately low relative to the size of the adrenal cortex. While most steroidogenic pathway genes downregulated on DEX recovered by 1-week post-withdrawal, *StAR* trended towards reduced expression relative to controls despite exposure to 2-fold higher ACTH levels. Post-withdrawal GIAI thus involves adrenocortical cell loss and a superimposed defect in steroidogenesis, which may be driven in part by impaired activation of the rate-limiting step in CORT production (Fig. S6).

Intermittent cosyntropin treatment partially attenuated DEX-induced adrenocortical apoptosis but paradoxically delayed post-withdrawal functional recovery. In contrast, CRH-driven, endogenous ACTH signaling during chronic DEX treatment preserved adrenal size, function, and morphology. Sustained trophic stimulation of the adrenal during chronic glucocorticoid exposure may thus represent a promising therapeutic strategy to prevent GIAI.

Our finding of rapid hypothalamic-pituitary recovery counters the longstanding assumption that prolonged central suppression limits HPA axis recovery after chronic glucocorticoid exposure. While prior rodent studies have also demonstrated rapid central axis recovery after glucocorticoid withdrawal^23–25^; we extended treatment to a duration known to cause protracted GIAI in humans, strengthening the translational relevance of this conclusion. These findings support repeating the clinical studies that established the central suppression paradigm using modern, direct ACTH assays with sampling timed to distinguish active glucocorticoid feedback from persistent hypothalamic pituitary dysfunction^14,16^.

The absence of central suppression in our model allowed us to investigate mechanisms of GIAI localized to the adrenal. Since ACTH is a well-established trophic hormone for the zF^7,8^, the time required for cellular regeneration from DEX-induced cortical atrophy offers an intuitive explanation for persistently low CORT secretion after glucocorticoid withdrawal despite central recovery.. The adult zona fasciculata is considered a moderately proliferative tissue with a turnover rate of 120 days in mice^48,49^. Although this exceeds the 8 weeks required for functional recovery in our study, accelerated renewal would be expected under conditions of increased ACTH secretion, as occurs after DEX withdrawal^50^. Interestingly, we found that adrenal mass recovered more rapidly than steroidogenic function. This is consistent with prior studies reporting low CORT levels and steroidogenic pathway gene expression one week after discontinuing short- to moderate-term glucocorticoids despite normalization of adrenal weights^23–25^. Using quantitative morphometry, we further demonstrated that this dissociation between adrenal size and function persisted even after accounting for the DEX-induced expansion of large, lipid-laden adrenal macrophages, suggesting that zF cell loss alone is insufficient to explain impaired CORT secretion after glucocorticoid withdrawal.

We therefore sought to identify a superimposed defect in steroidogenesis and found that *Star* expression remained below control levels in DEX-withdrawn adrenals despite supraphysiologic ACTH stimulation. StAR acutely stimulates steroidogenesis by facilitating cholesterol transfer to the inner mitochondrial membrane, thereby making substrate available to CYP11A1 for the first step in steroid hormone synthesis^51–53^. Prior rodent studies have reported decreased *Star* expression 5-7 days after discontinuing short- to moderate-term glucocorticoid treatment, though these studies utilized bulk methodologies potentially confounded by decreased zF cell representation in adrenal homogenates^23,25^. RNAScope analysis localized this pattern to individual zF cells, consistent with a steroidogenic defect superimposed on zF atrophy. We also evaluated *Star* expression following insulin-induced hypoglycemia rather than at rest, since *Star* is rapidly transcribed on demand via ACTH-dependent cAMP/PKA signaling in response to physiologic stress^54^. The apparent failure of *Star* induction despite two-fold higher circulating ACTH concentrations contrasted with the response of *Hsd3b2*, which is also regulated by ACTH/cAMP/PKA signaling but trended toward 2-fold higher expression relative to controls in the same DEX-withdrawn animals.

It is important to note that these expression differences in both *Star* and *Hsd3b2* in 8+1 vs. DN adrenals did not reach statistical significance (p=0.12 and 0.13, respectively) and should be treated as hypothesis-generating until larger, adequately powered studies can be performed.

The selective failure to induce *Star* during hypoglycemic stress is nonetheless of mechanistic interest in light of a paracrine signaling mechanism proposed by Xu *et al.,* in which activated adrenal macrophages suppress *Star* in neighboring zF cells^43^. In this model, rising intra-adrenal glucocorticoids during stress upregulate *Trem2* in adrenal macrophages, triggering TGF-β secretion and downstream StAR suppression in the zF^43^. Consistent with this mechanism, the large, lipid-laden macrophages in our DEX-withdrawn adrenals robustly expressed *Trem2* during hypoglycemic stress. Future studies will determine whether adrenal macrophage depletion or disruption of this pathway restores *Star* expression and CORT secretion.

Adrenal macrophage accumulation has not previously been reported after chronic glucocorticoid treatment, though Wylie *et al* detected apoptotic bodies “within phagosomes of histiocytes” during short-term prednisolone exposure^7^. Like macrophages described in acute and chronic stress states, those in our model express classical macrophage markers (CD68, F4/80, CD11c), are long-lived, and are intensely autofluorescent due to lipid accumulation^43,55–57^. They most strikingly differ from typical adrenal macrophages in both their large size and apparent multinucleation; this may reflect macrophage fusion under conditions of high phagocytic demand, as has been described in adipose, ovarian, and bone tissues^58–60^.

Morphologically similar macrophages have been observed in aged, male *Znrf3* knockout mice, which undergo early adrenocortical hyperplasia followed by senescence^61^, suggesting this phenotype is not unique to glucocorticoid exposure but may instead be a feature of adrenal remodeling after cortical cell loss.

Most importantly, identifying the adrenal as the rate-limiting step in HPA axis recovery opens it as an accessible therapeutic target for GIAI prevention that resides outside the blood-brain barrier. We have shown that sustained, CRH driven ACTH signaling, but not intermittent cosyntropin treatment, preserves adrenal function during chronic glucocorticoid exposure. The failure of cosyntropin to prevent GIAI in our model aligns with a prior clinical study in which combined intramuscular ACTH and glucocorticoid treatment produced more severe HPA axis suppression than glucocorticoids alone^62^. Both ACTH and cosyntropin have extremely short half-lives of 10-30 minutes^63,64^, and even administration of cosyntropin at doses more than 100-fold above the physiologic circadian peak, as done in this study, cannot replicate the pulsatile secretory pattern of endogenous ACTH^65^. While the functional importance of ACTH pulsatility is incompletely understood, pulsatile but not continuous ACTH infusion restored CORT secretion and steroidogenic enzyme gene expression in glucocorticoid-treated rats^66,67^. Ultradian ACTH dynamics could not be assessed in HGRKO mice due to blood volume constraints and the stress of serial sampling, though circadian ACTH and CORT rhythms are preserved^27^. DEX-treated HGRKO mice also likely continue to secrete other POMC-derived peptides known to stimulate adrenal mitogenesis, which would be absent in cosyntropin-rescued WT animals^68,69^.

Consistent with the conclusion that intermittent cosyntropin did not reproduce physiologic ACTH signaling, *Mc2r* expression was equally suppressed in short-term DEX- and DEX+COS-treated WT animals. Surprisingly, per-cell *Mc2r* expression was also reduced in DEX-treated HGRKO mice relative to controls. ACTH regulates MC2R in a biphasic fashion, with moderate levels stimulating transcription but higher levels promoting receptor desensitization^70,71^. This latter negative feedback mechanism may account for the lower per-cell expression observed in HGRKO mice relative to DN controls despite sustained ACTH exposure. Single-cell *Mc2r* mRNA content was also unexpectedly similar in the small population of zF cells that survived long-term DEX-treatment (8+0) relative to controls. This may reflect either selection for a zF subpopulation less dependent on ACTH signaling or impaired adrenal zonation in the setting of chronic ACTH deficiency giving rise to cells with mixed zF and zG characteristics^48,50,72,73^.

While even sustained ACTH-based therapeutic strategies may therefore be of limited utility in preserving adrenal function during chronic glucocorticoid treatment, our findings in HGRKO mice highlight the promise of long-acting CRH analogs as a strategy to prevent GIAI. In patients with Cushing syndrome, adrenal function recovers more rapidly after resection of an ectopic, CRH-producing tumor than an ACTH-secreting pituitary adenoma^74^. GR expression is intact in the ACTH-positive corticotrophs of the anterior pituitary in HGRKO mice^27^, indicating that CRH stimulation can dominate over glucocorticoid-mediated inhibition at the level of the pituitary. The pituitary’s location outside the blood-brain barrier further supports the translational feasibility of systemically administered CRH-based therapeutics. We have previously demonstrated that continuous subcutaneous infusion of CRH restores circadian ACTH and corticosterone rhythms in CRH-knockout mice^32^. Future work will determine whether systemic CRH administration during long-term glucocorticoid treatment phenocopies the HGRKO mouse by producing physiologic ACTH secretion that preserves adrenal function, and critically, whether these effects are sustained after both CRH and glucocorticoid therapy are discontinued.”

As a potential limitation of the HGRKO model, its endocrine phenotype arises from hypothalamic GR deletion, which disrupts the principal mechanism by which glucocorticoids exert negative feedback on the HPA axis. Accordingly, the HGRKO does not provide a directly translatable therapy for GIAI but instead serves as an important proof of principle that maintenance of physiologic trophic support to the adrenal during long-term glucocorticoid therapy preserves CORT output. The constitutive nature of this model precluded assessment of whether full HPA axis function is restored following both glucocorticoid withdrawal and reversal of the genetic manipulation. Although this will be an important consideration for future attempts to replicate this model pharmacologically, we do not anticipate it to represent a major limitation given our finding that hypothalamic-pituitary suppression reverses within one week of DEX withdrawal even in wild-type mice.

An additional limitation is that our study included only male mice to minimize biological variability across a broad, exploratory survey of HPA axis recovery. While there are no reported sex differences in the risk, severity, or duration of GIAI in humans^10,75–77^, HPA axis responsiveness, rates of adrenocortical turnover, and adrenal macrophage populations are all sexually dimorphic in rodents^61,78–84^. Future studies will specifically investigate sex differences in the mechanisms of post-withdrawal GIAI and time to HPA axis recovery in this preclinical model. Additionally, sample sizes were relatively modest to permit interrogation of HPA axis function at multiple timepoints after chronic glucocorticoid treatment. Since we examined HPA axis function in the stimulated state, as is done to clinically diagnose adrenal insufficiency, analyses of all primary hormonal outcomes were all still adequately powered. Nonetheless, subsequent mechanistic studies will include larger cohorts of animals now that biologically relevant timepoints have been identified.

In conclusion, our findings demonstrate that post-withdrawal GIAI arises primarily from adrenal dysfunction rather than delayed hypothalamic-pituitary recovery. This prolonged adrenal suppression, driven by both adrenocortical cell loss and impaired steroidogenesis, is prevented by maintaining sustained trophic stimulation of the adrenal during glucocorticoid exposure.

These results establish the adrenal as the principal locus of post-withdrawal GIAI and an accessible target for therapeutic intervention. Pharmacologic strategies that maintain physiologic adrenal stimulation could prevent GIAI, thereby reducing the morbidity associated with one of the most widely prescribed drug classes while preserving their therapeutic efficacy.

## Supporting information

Supplemental Figures S1-S6 and Tables SI-SIII

## Acknowledgments

We thank Drs. David Breault, Kleiton Borges, and Mesut Berber for their input on this project and assistance processing adrenal tissue, identifying zonal markers, and performing thick, floating section immunofluorescence. We also thank Suzanne White from the BIDMC Histology core for preparing and sectioning all FFPE specimens, including working closely with the author (LG) to isolate the PVH in mouse brains.

## Data availability

Original data generated and analyzed during this study are included in this published article or in the data repositories listed in References^28^.

This study was supported by NIH 5T32DK007699-39 (LG), NIH F32DK131795-02 (LG), NIH 1K08DK142004-01A1 (LG), a Pediatric Endocrine Society Research Fellowship Award (LG), a Pediatric Endocrine Society Clinical Scholar Award (LG), a Boston Children’s Hospital Office of Faculty Development/Basic & Clinical Translational Research Executive Committees Faculty Career Development Fellowship (LG), and the Thomas Morgan Rotch Harvard Medical School endowment (JAM).

