## Supplemental Figures S1-S6 and Tables SI-SIII for "Adrenal rather than central dysfunction limits HPA axis recovery after chronic glucocorticoid treatment in male mice"

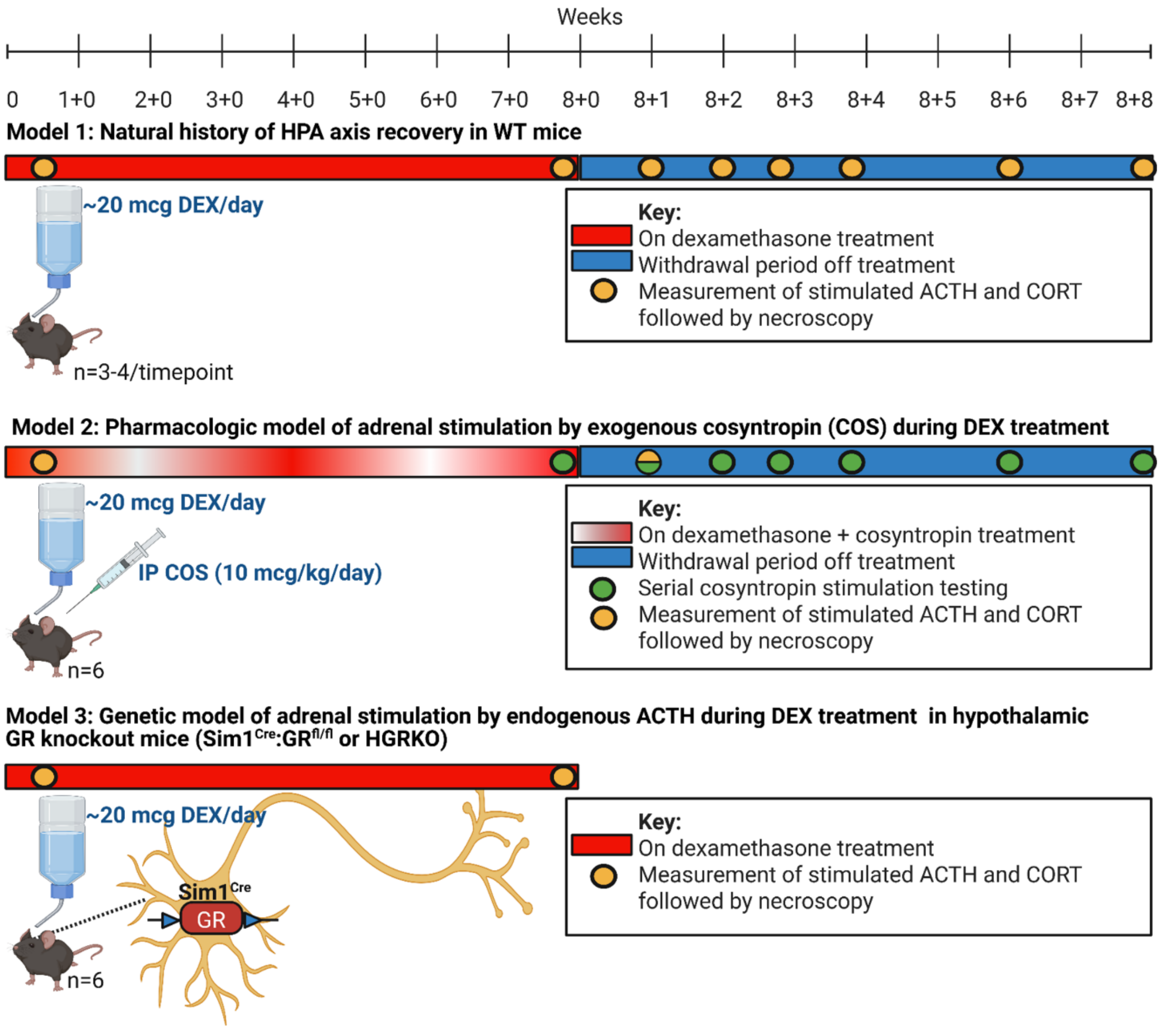


**Figure S1. Experimental design for dexamethasone (DEX) ± cosyntropin (COS) treatment and withdrawal in wild-type (WT; Models 1 and 2) and *Sim1*$^{Cre}$:*GR*$^{fl/fl}$ (HGRKO; Model 3) mice.** In model 1, the natural history of HPA axis recovery after long-term DEX was studied by treating WT mice with 8 weeks of DEX (red bar) followed by serial assessments of axis function and histology at 1-2 week intervals (yellow dots) after DEX was withdrawn (blue bar). Insulin-induced hypoglycemia and cosyntropin stimulation testing were performed on successive days followed by necroscopy. Pharmacologic (model 2) and genetic strategies (model 3) were then studied for their ability to prevent GIAI. In model 2, mice were co-treated for 8 weeks with DEX and cosyntropin, an ACTH analog, followed by cosyntropin-stimulation testing to assess adrenal function after both agents were withdrawn (green dots). Necroscopy for histologic studies was performed in a separate cohort of mice 1-week post-DEX+COS (8+1; split yellow and green dot). In model 3, HPA axis activity was assessed upon completion of an 8-week DEX course (8+0) in mice with selective hypothalamic deletion of the glucocorticoid receptor (HGRKO), which allows for ongoing, endogenous ACTH secretion on GC therapy. ACTH and CORT were measured in the basal, unstimulated state followed by provocative assessments (restraint stress and cosyntropin-stimulation testing on successive days) and necroscopy. Created in BioRender. Gaston, L. (2026) https://BioRender.com/u7gkjwg.

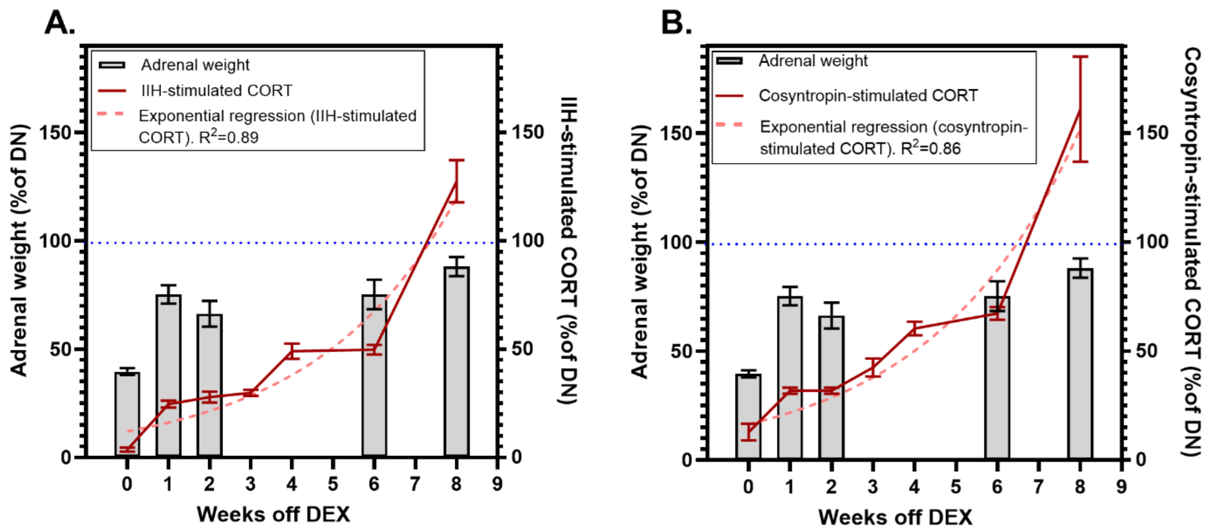


**Figure S2. Adrenal mass and stimulated CORT secretion follow different recovery trajectories after long-term DEX exposure.** Adrenal weights (left Y-axis, gray bars) and A) insulin-induced hypoglycemia (IIH)-stimulated- or B) cosyntropin-stimulated CORT secretion (right Y-axis, red solid line) vs. weeks after withdrawal of 8 weeks of DEX treatment (X-axis). Adrenal weight data in A and B are identical and derived from those in Fig. 2*A* as a percentage of DN controls. CORT data are derived from those in Figs. *1D* and *E* as a percentage of their DN controls. Pink dashed lines denote exponential growth models for IIH-stimulated ($Y = 12.1 * e^{0.29X}$, doubling time = 2.4 weeks, $R^2$=0.89) and cosyntropin-stimulated ($Y = 16.5 * e^{0.28X}$, doubling time = 2.5 weeks, $R^2$=0.86) CORT secretion.

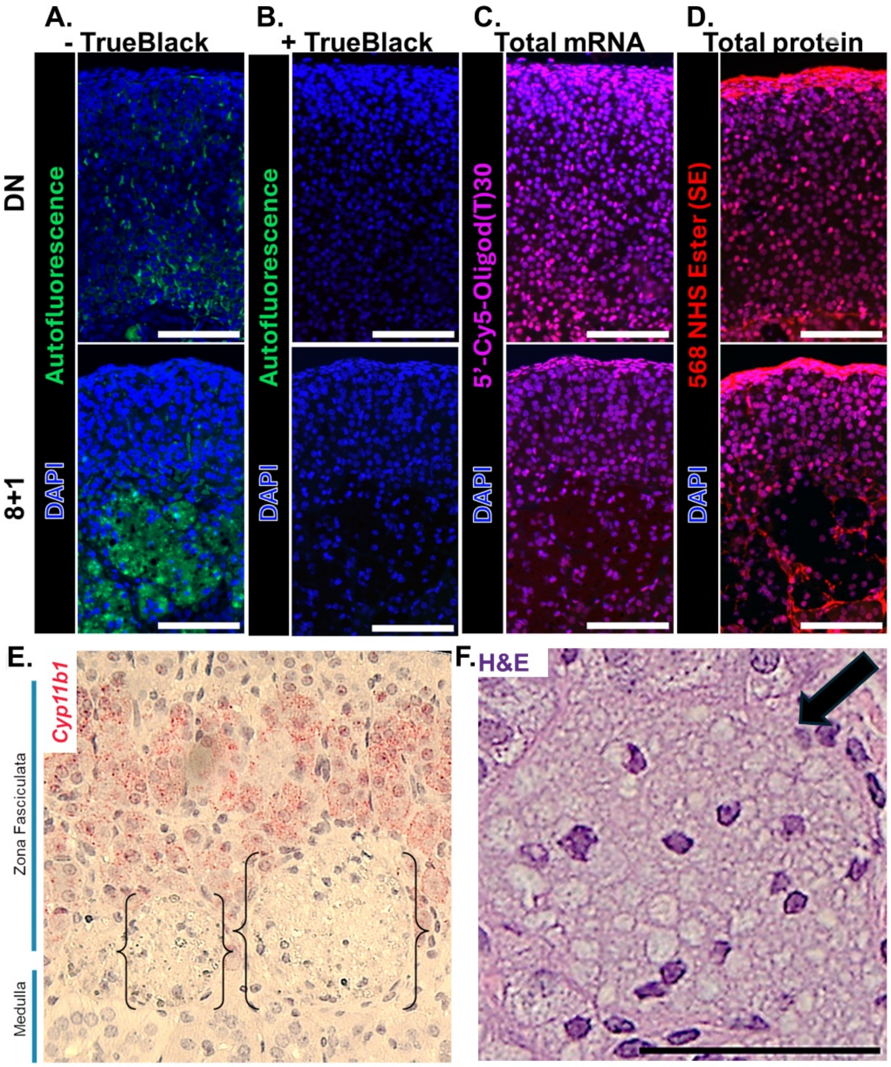


**Figure S3. The large, vacuolated structures that accumulate during long-term DEX treatment contain lipofuscin-rich, transcriptionally active cells rather than proteinaceous debris but are distinct from surrounding zona fasciculata cells.** A) DEX-exposed adrenals (8+1, bottom) contain intensely autofluorescent clusters (green) that are absent in DN controls (top). B) Autofluorescence is eliminated with the addition of TrueBlack, a lipofuscin autofluorescence quencher. Nuclei within these clusters contain C) mature mRNA (*in situ* hybridization for total polyA with 5'-Cy5-Oligo d(T)30, magenta; with TrueBlack). D) The surrounding area is not intensely labeled by 568 NHS Ester (red; with TrueBlack), a total protein stain. Scale bars=100 microns. E) Cells in these clusters (bracketed) do not express the steroidogenic enzyme *Cyp11b1*, a zF marker (red, *in situ* hybridization using RNAScope with hematoxylin nuclear counterstain in a representative 8+1 adrenal). F) Enlarged H&E image of a representative macrophage (black arrow) in an 8+0 adrenal taken at 20x magnification. Scale bar=50 microns.

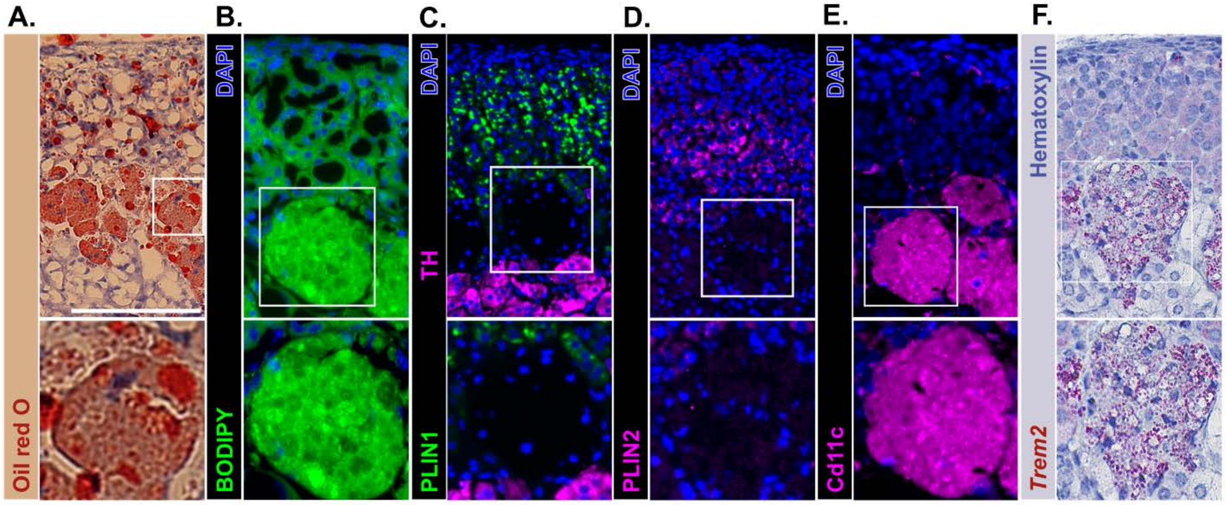


**Figure S4. Adrenal macrophages in DEX-exposed adrenals are lipid-filled, negative for lipid droplet associated proteins perilipin-1 and 2, and express some genes characteristic of lipid-associated macrophages.** For all stains, representative images shown from 8+1 adrenals (top) with magnification of a macrophage cluster (bottom). A) Oil red O and B) BODIPY 493/503 staining. Immunofluorescence for C) perilipin-1 (PLIN1; green) with tyrosine hydroxylase (TH; magenta) to label the adrenal medulla, D) perilipin-2 (PLIN2; magenta), and E) Cd11c (magenta). F) *In situ hybridization* (RNAScope) for *Trem2* (red) with hematoxylin nuclear counterstain. Scale bar=100 microns.

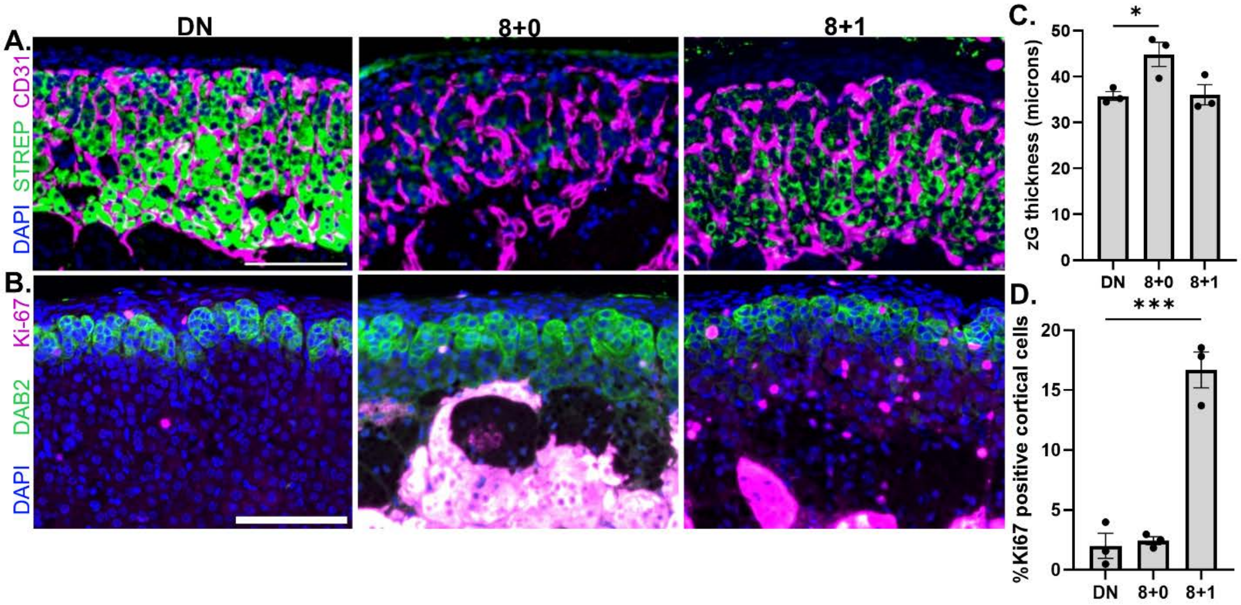


**Figure. S5. DEX-withdrawn adrenals have an intact vascular network, normal zG thickness, and a marked increase in zF proliferation.** Immunofluorescence for A) for the vascular marker CD31 (magenta) with a 488-streptavidin counterstain (green) to mark steroidogenic cortical cells and B) the proliferative marker Ki-67 (magenta) with Dab2 (green) to define the zG. Representative images from DN, 8+0, and 8+1 glands with quantification of C) zG thickness ($p=0.03$ by one-way ANOVA), and D) the percentage of Ki-67 positive adrenocortical cells ($p=0.0001$). $^*p<0.05$ and $^{***}p<0.001$ for post-hoc pairwise comparisons vs. DN with Bonferroni correction. Scale bar=100 microns.

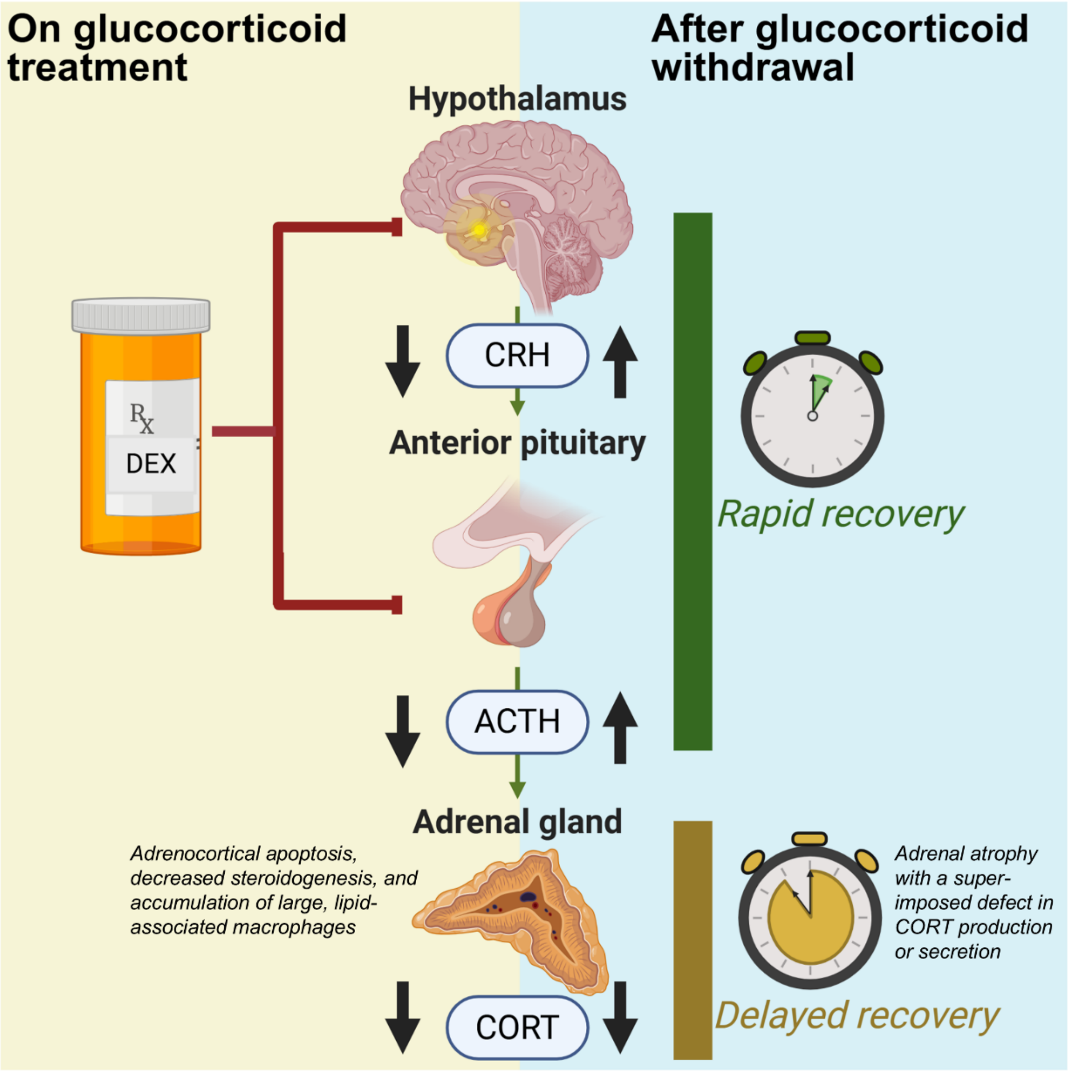


**Figure S6. Adrenal rather than hypothalamic-pituitary dysfunction limits HPA axis recovery after chronic glucocorticoid treatment in male mice**. Schematic of the proposed mechanism underlying prolonged GIAI following steroid discontinuation. During chronic glucocorticoid treatment (left; yellow), central glucocorticoid-mediated negative feedback suppresses CRH and ACTH secretion from the hypothalamus and anterior pituitary, respectively. Under conditions of reduced ACTH stimulation, adrenal zona fasciculata cells undergo apoptosis with concomitant suppression of steroidogenesis, and the gland becomes progressively infiltrated by lipid-laden macrophages. Upon withdrawal of chronic glucocorticoid treatment (right; blue), hypothalamic-pituitary activity is rapidly restored; however, adrenal insufficiency persists for months, driven by both adrenocortical cell loss and a superimposed defect in CORT production or secretion.

**Table SI. PCR primers and settings for genotyping $Sim1^{Cre}$:$GR^{fl/fl}$ mice and controls (HGRKO and HGRWT).**

| Genotype reaction | Primer sequence | PCR settings |
|---|---|---|
| GR FLOX | Forward: ATG CCT GCT AGG CAA ATG AT<br>Reverse: TTC CAG GGC TAT AGG AAG CA | Stage 1: 95°C 3 min (1x);<br>Stage 2: 95°C 30 sec - 56°C 30 sec – 52°C 30 sec (10x);<br>Stage 3: 68°C 1 min (1x);<br>Stage 4: 95°C 30 sec - 53°C 30 sec - 68°C 2 min (35x);<br>Stage 5: 68°C 2min (1x)<br>4°C hold |
| Common Cre | Forward: TGC ATG ATC TCC GGT ATT GA<br>Reverse: CGT ACT GAC GGT GGG AGA AT | Stage 1: 94°C 2 min (1x);<br>Stage 2: 94°C 30 sec - 55°C 30 sec - 72°C 1 min (30x)<br>Stage 3: 72°C 5 min (1x)<br>4°C hold |

**Table SII. Antibodies used for paraffin section immunofluorescence**

| Antibody | Antigen retrieval | Dilution | Company | Catalog number | RRID |
|---|---|---|---|---|---|
| *Primary antibodies* | | | | | |
| Rabbit anti-CD68 | Sodium Citrate pH 6.0 | 1:750 | Abcam | ab125212 | AB_10975465 |
| Rabbit anti-F4/80 | Tris-EDTA pH 9.0 | 1:100 | CST | 70076T | AB_2799771 |
| Rabbit anti-Perilipin-1 | Sodium Citrate pH 6.0 | 1:100 | CST | 9349 | AB_10829911 |
| Rabbit anti-Perilipin-2 | Tris-EDTA pH 9.0 | 1:200 | CST | 95109 | AB_3711338 |
| Rabbit anti-Cd11c | Tris-EDTA pH 9.0 | 1:300 | CST | 97585 | AB_2800282 |
| Sheep anti-tyrosine hydroxylase | Tris-EDTA pH 9.0 | 1:5000 | Sigma Aldrich | AB1542 | AB_90755 |
| Rabbit anti-CD31 | Tris-EDTA pH 9.0 | 1:100 | CST | 77699 | AB_2722705 |
| Rabbit anti-Ki-67 | Tris-EDTA pH 9.0 | 1:100 | CST | D3B5 | AB_2922798 |
| Mouse (IgG1) anti-Dab2 | Tris-EDTA pH 9.0 | 1:200 | BD Biosciences | 610464 | AB_397837 |
| *Secondary antibodies* | | | | | |
| Donkey anti-rabbit IgG/647 | | 1:200 | Invitrogen | A-31573 | AB_2536183 |
| Donkey anti-rabbit IgG/488 | | 1:200 | Invitrogen | A-21206 | AB_2535792 |
| Goat anti-mouse IgG1/568 | | 1:200 | Invitrogen | A-21124 | AB_2535766 |
| Donkey anti-sheep IgG/647 | | 1:200 | Invitrogen | A-21448 | AB_2535865 |

**Table SIII. Normality and equal variance testing to guide statistical test selections.** Shapiro-Wilk test for normality and Brown-Forsythe test for homogeneity of variance were applied to all outcome variables. Parametric tests (Welch's t-test, one-way ANOVA with Bonferroni) correction were used when both assumptions were satisfied; non-parametric alternatives (Mann-Whitney U, Kruskal-Wallis with Dunn's correction) were used otherwise.

| Variable | Figure | Group | n | Shapiro-Wilk W | p | Brown-Forsythe F (DFn, DFd) | p |
|---|---|---|---|---|---|---|---|
| PVH *Crh* expression | 1B | | | | | 0.2837 (2, 6) | 0.763 |
| | | DN | 3 | 0.7997 | 0.1137 | | |
| | | 8+0 | 3 | 0.9573 | 0.6023 | | |
| | | 8+1 | 3 | 0.9071 | 0.4086 | | |
| IIH-stimulated ACTH | 1C | | | | | 1.476 (6, 18) | 0.242 |
| | | DN | 4 | 0.8691 | 0.2941 | | |
| | | 8+0 | 4 | 0.706 | 0.0136[†] | | |
| | | 8+1 | 4 | 0.8666 | 0.2847 | | |
| | | 8+2 | 3 | 0.9226 | 0.4616 | | |
| | | 8+3 | 4 | 0.907 | 0.4668 | | |
| | | 8+4 | 3 | 0.9996 | 0.9612 | | |
| | | 8+6 | 4 | 0.8514 | 0.2308 | | |
| | | 8+8 | 3 | 0.9765 | 0.706 | | |
| IIH-stimulated CORT | 1D | | | | | 0.6908 (7, 21) | 0.679 |
| | | DN | 4 | 0.8928 | 0.3962 | | |
| | | 8+0 | 4 | 0.9955 | 0.9838 | | |
| | | 8+1 | 4 | 0.9515 | 0.7256 | | |
| | | 8+2 | 3 | 0.9868 | 0.7804 | | |
| | | 8+3 | 4 | 0.9399 | 0.6536 | | |
| | | 8+4 | 3 | 0.9113 | 0.4226 | | |
| | | 8+6 | 4 | 0.8997 | 0.4296 | | |
| | | 8+8 | 3 | 0.7796 | 0.0668 | | |
| Cosyntropin-stimulated CORT | 1E | | | | | 2.044 (7, 20) | 0.099 |
| | | DN | 4 | 0.9768 | 0.883 | | |
| | | 8+0 | 4 | 0.9608 | 0.7841 | | |
| | | 8+1 | 3 | 0.8538 | 0.2508 | | |
| | | 8+2 | 3 | 0.9745 | 0.6937 | | |
| | | 8+3 | 4 | 0.9758 | 0.8767 | | |
| | | 8+4 | 3 | 0.8578 | 0.2615 | | |
| | | 8+6 | 4 | 0.9202 | 0.5381 | | |
| | | 8+8 | 3 | 0.9464 | 0.5537 | | |
| Adrenal weight | 2A | | | | | 0.8081 (5, 18) | 0.559 |
| | | DN | 4 | 0.8949 | 0.4064 | | |
| | | 8+0 | 4 | 0.9447 | 0.683 | | |

| | | | | | | | |
|---|---|---|---|---|---|---|---|
| | | 8+1 | 4 | 0.9271 | 0.5774 | | |
| | | 8+2 | 4 | 0.989 | 0.9523 | | |
| | | 8+6 | 4 | 0.8397 | 0.1945 | | |
| | | 8+8 | 4 | 0.9631 | 0.7982 | | |
| Pan-cortical area | 3B | | | | | N/A[a] | |
| | | DN | 3 | 0.9453 | 0.5492 | | |
| | | 8+0 | 3 | 0.7617 | 0.026 | | |
| | | 8+1 | 3 | 0.9991 | 0.9424 | | |
| | | 8+2 | 3 | 0.7576 | 0.0168 | | |
| | | 8+6 | 2 | Unable to calculate; excluded from analysis | | | |
| | | 8+8 | 3 | 0.9322 | 0.497 | | |
| Medullary area | 3C | | | | | N/A[a] | |
| | | DN | 3 | 0.9491 | 0.5656 | | |
| | | 8+0 | 3 | 0.8397 | 0.2134 | | |
| | | 8+1 | 3 | 0.7586 | 0.0191 | | |
| | | 8+2 | 3 | 0.9965 | 0.8865 | | |
| | | 8+6 | 2 | Unable to calculate; excluded from analysis | | | |
| | | 8+8 | 3 | 0.9993 | 0.9504 | | |
| Adrenal macrophage area | 3D | DN | 3 | 0.8357 | 0.2029 | N/A[a] | |
| | | 8+0 | 3 | 0.8434 | 0.2229 | | |
| | | 8+1 | 3 | 0.9406 | 0.5298 | | |
| | | 8+2 | 3 | 0.8632 | 0.2763 | | |
| | | 8+6 | 2 | Unable to calculate; excluded from analysis | | | |
| | | 8+8 | 3 | 0.7578 | 0.0173 | | |
| Steroidogenic area | 3E | | | | | 0.9073 (5, 11) | 0.51 |
| | | DN | 3 | 0.942 | 0.5354 | | |
| | | 8+0 | 3 | 0.9525 | 0.5805 | | |
| | | 8+1 | 3 | 0.9985 | 0.9249 | | |
| | | 8+2 | 3 | 0.8266 | 0.1796 | | |
| | | 8+6 | 2 | Unable to calculate; excluded from analysis | | | |
| | | 8+8 | 3 | 0.9671 | 0.6518 | | |
| Normalized stimulated CORT/steroidogenic area | 3F | | | | | 1.124 (5, 11) | 0.403 |
| | | DN | 3 | 0.9948 | 0.8621 | | |

| | | | | | | | |
|---|---|---|---|---|---|---|---|
| | | 8+0 | 3 | 0.9999 | 0.9769 | | |
| | | 8+1 | 3 | 0.9973 | 0.9003 | | |
| | | 8+2 | 3 | 0.8563 | 0.2575 | | |
| | | 8+6 | 2 | Unable to calculate; excluded from analysis | | | |
| | | 8+8 | 3 | 0.9899 | 0.8078 | | |
| zF *Mc2r* expression | 4F | | | | | N/A[a] | |
| | | DN | 4 | 0.8674 | 0.2878 | | |
| | | 8+0 | 3 | 0.7651 | 0.0338 | | |
| | | 8+1 | 5 | 0.9175 | 0.5137 | | |
| zF *Star* expression | 4G | DN | 4 | 0.8882 | 0.3747 | 0.8565 (2, 9) | 0.457 |
| | | 8+0 | 3 | 0.9861 | 0.7745 | | |
| | | 8+1 | 5 | 0.8547 | 0.2098 | | |
| zF *Cyp11a1* expression | 4H | | | | 0.9076 | N/A[a] | |
| | | DN | 4 | 0.9809 | | | |
| | | 8+0 | 3 | | 0.0155 | | |
| | | 8+1 | 5 | 0.9254 | 0.5651 | | |
| zF *Hsd3b* expression | 4I | | | | | | |
| | | DN | 4 | 0.8855 | 0.3628 | 1.373 (2, 9) | 0.302 |
| | | 8+0 | 3 | 0.9516 | 0.5765 | | |
| | | 8+1 | 5 | 0.8669 | 0.254 | | |
| zF *Cyp11b1* expression | 4J | | | | | 1.161 (2, 9) | 0.356 |
| | | DN | 4 | 0.8853 | 0.3618 | | |
| | | 8+0 | 3 | 0.9865 | 0.7777 | | |
| | | 8+1 | 5 | 0.963 | 0.829 | | |
| %TUNEL positive cells | 5B | | | | | 6.796 (2, 13) | 0.01 |
| | | DN | 5 | 0.9664 | 0.8516 | | |
| | | 4D DEX | 4 | 0.9792 | 0.8972 | | |
| | | 4D DEX+COS | 7 | 0.8867 | 0.2576 | | |
| DEX + COS cosyntropin-stimulated CORT | 5C | 8+0 | 6 | 0.7906 | 0.0483 | N/A[b] | |
| | | 8+1 | 6 | 0.9217 | 0.5177 | | |
| | | 8+2 | 5 | 0.9514 | 0.7468 | | |
| | | 8+4 | 5 | 0.9123 | 0.4813 | | |
| | | 8+6 | 5 | 0.8758 | 0.2906 | | |
| | | 8+8 | 5 | 0.9085 | 0.4589 | | |
| Pan-cortical area (normalized to DN) | 5E | 8+1 | 3 | 0.9991 | 0.9424 | N/A[c] | |
| | | DEX+COS 8+1 | 3 | 0.8536 | 0.2501 | | |

| | | | | | | | |
|---|---|---|---|---|---|---|---|
| Adrenal macrophage area (normalized to DN) | 5F | 8+1 | 3 | 0.9406 | 0.5298 | N/A[d] | |
| | | DEX+COS 8+1 | 3 | 0.7542 | 0.0094 | | |
| Steroidogenic cell area (normalized to DN) | 5G | 8+1 | 3 | 0.9985 | 0.9249 | N/A[c] | |
| | | DEX+COS 8+1 | 3 | 0.8411 | 0.217 | | |
| %TUNEL positive cells | 7B | DN | 5 | 0.9664 | 0.8516 | 8.041 (2, 10) | 0.008 |
| | | 4D DEX | 4 | 0.9792 | 0.8972 | | |
| | | 4D HGRKO | 4 | 0.8116 | 0.1248 | | |
| Basal ACTH | 7C | | | | | | |
| | | Pre-DEX: HGRWT | 3 | 0.9647 | 0.6389 | N/A[c] | |
| | | Pre-DEX: HGRKO | 5 | 0.8942 | 0.3784 | | |
| | | On 8W DEX: HGRWT | 3 | Unable to calculate | | N/A[d] | |
| | | On 8W DEX: HGRKO | 6 | 0.9472 | 0.7179 | | |
| CORT | 7D | Pre-DEX: HGRWT, Basal | 3 | 0.7711 | 0.0472 | N/A[d] | |
| | | Pre-DEX: HGRKO, Basal | 6 | 0.9128 | 0.455 | | |
| | | Pre-DEX: HGRWT, Restraint | 4 | 0.9076 | 0.47 | N/A[c] | |
| | | Pre-DEX: HGRKO, Restraint | 4 | 0.9137 | 0.5023 | | |
| | | On 8W DEX: HGRWT, Basal | 3 | 0.9353 | 0.5087 | N/A[c] | |
| | | On 8W DEX: HGRKO, Basal | 6 | 0.9225 | 0.5232 | | |
| | | On 8W DEX: HGRWT, Restraint | 3 | 0.8311 | 0.1911 | N/A[c] | |
| | | On 8W DEX: HGRKO, Restraint | 6 | 0.9296 | 0.5767 | | |
| | | On 8W DEX: HGRWT, COS-stim | 3 | 0.7547 | 0.0103 | N/A[d] | |
| | | On 8W DEX: HGRKO, COS-stim | 6 | 0.9624 | 0.8384 | | |
| Adrenal weight | 7E | HGRWT | 4 | 0.7286 | 0.0239 | N/A[d] | |
| | | HGRKO | 4 | 0.9447 | 0.683 | | |
| Pan-cortical area (normalized to DN) | 7F | HGRWT 8+0 | 3 | 0.9047 | 0.4006 | N/A[c] | |
| | | HGRKO 8+0 | 3 | 0.7827 | 0.0738 | | |
| AM area (normalized to DN) | 7G | HGRWT 8+0 | 3 | 0.7781 | 0.0631 | N/A[c] | |

| | | | | | | | |
|---|---|---|---|---|---|---|---|
| | | HGRKO 8+0 | 3 | 0.8729 | 0.3038 | | |
| Steroidogenic cell area (normalized to DN) | 7H | HGRWT 8+0 | 3 | 0.8089 | 0.1358 | N/A[c] | |
| | | HGRKO 8+0 | 3 | 0.815 | 0.1509 | | |
| zF *Mc2r* expression (AU) | 8B | | | | | 1.466 (3, 12) | 0.273 |
| | | DN | 4 | 0.8868 | 0.3684 | | |
| | | WT 4D DEX | 4 | 0.9104 | 0.4845 | | |
| | | 4D DEX+COS | 4 | 0.8906 | 0.386 | | |
| | | HGRKO 4D DEX | 4 | 0.839 | 0.1926 | | |
| zG thickness | S5C | DN | 3 | 0.9137 | 0.4305 | 0.2604 (2, 6) | 0.779 |
| | | 8+0 | 3 | 0.8585 | 0.2635 | | |
| | | 8+1 | 3 | 0.8203 | 0.1639 | | |
| %Ki67-positive cortical cells | 5D | DN | 3 | 0.9535 | 0.5851 | 0.5461 (2, 6) | 0.606 |
| | | 8+0 | 3 | 0.9861 | 0.7746 | | |
| | | 8+1 | 3 | 0.8599 | 0.2673 | | |

†:Hormone concentrations in the DEX-treated group were non-normally distributed, with multiple values below the assay's lower limit of detection; this group was therefore excluded from post-withdrawal comparisons, which satisfied assumptions for parametric analysis.

a: Multi-group comparisons in which at least one group failed to satisfy normality were performed using the Kruskal-Wallis test, a non-parametric alternative to one-way ANOVA that does not require normality or homogeneity of variance. Pairwise post-hoc comparisons were performed using Dunn's test with correction for multiple comparisons.

b: Homoscedasticity of regression residuals was confirmed by visual inspection of a residual plot (absolute residuals vs. fitted values), which showed no systematic relationship between residual variance and fitted values.

c: Two-group comparisons satisfying normality were performed using Welch's unpaired t-test, which applies a correction for unequal variances and therefore does not require prior assessment of homoscedasticity.

d: Two-group comparisons in which at least one group failed to satisfy normality were performed using the Mann-Whitney U test, a non-parametric alternative that does not require normality or prior assessment of homoscedasticity.
